# Genetical engineered lung cancer cell for analyzing Epithelial-Mesenchymal transition

**DOI:** 10.1101/778316

**Authors:** Michał Kiełbus, Jakub Czapiński, Joanna Kałafut, Justyna Woś, Andrzej Stepulak, Adolfo Rivero-Müller

## Abstract

Cell plasticity, defined as the ability to undergo phenotypical transformation in a reversible manner, is a physiological processes that also exert important roles in disease progression Two forms of cellular plasticity are epithelial-mesenchymal transition (EMT) and its inverse process, mesenchymal-epithelial transition (MET). These processes have been correlated to the poor outcome of different types of neoplasias as well as drug resistance development. Since EMT/MET are transitional processes, we have generated and validated a reporter cell line. Specifically, a far-red fluorescent protein was knocked-in in-frame with the mesenchymal gene marker *VIMENTIN* (*VIM*) in H2170 lung cancer cells. The vimentin reporter cells (VRCs) are a reliable model for studying EMT and MET showing cellular plasticity upon a series of stimulations. These cells are a robust platform to dissect the molecular mechanisms of these processes, and for drug discovery *in vitro* and in the future *in vivo*.

## 1. Introduction

The ability of cells to temporally acquire different characteristics, also known as cell plasticity, plays essential roles in physiological and pathophysiological processes [1]. Epithelial cells might transform to a mesenchymal phenotype and return to epithelial phenotype by epithelial-mesenchymal transition (EMT) and mesenchymal-epithelial transition (MET), respectively. EMT has been correlated to the metastasis and invasion potential of many types of malignancies [2]. The fact that the poor outcome of many types of neoplasias correlates with EMT/MET, makes these molecular phenomena important focus for research and drug targeting [3–6]. Despite the clinical association, the role of EMT/MET in metastasis is inconclusive, for example mesenchymal-like prostate cancer cells survive in circulation but, unlike epithelial or cells undergoing EMT, they are unable to form macrometastases [7]. In fact, breast cancer cell metastasis to lung tissue in mice was not affected by decreasing EMT - by targeting EMT-triggering transcription factors such as *SNAI1* (*Snail*) and *SNAI2* (*Slug*) by overexpressing miR-200 [8]. An additional role of EMT in tumorigenesis might be the development of apoptotic tolerance and increased resistance to chemotherapy as has been found in animal models and patients [8,9]. Recently, a hybrid stage between epithelial and mesenchymal phenotypes (hybE/M) has been recognized, and such hybE/M cells migrate outside the primary tumors displaying some mesenchymal features such as spindle-like morphology, increased nuclear levels of ZEB1 transcription factor, and epithelial ones such as cell-cell adhesion potential [10,11].

In order to study the transitory stages of EMT, MET and hybE/M, in particular at *in vivo* settings, there is need to generate reporter cells to visualize phenotypical changes. There have been different approaches to this, for example the use of heterologous fluorescent proteins driven under stage-specific promoters such as mesenchymal (*ZEB1*) and epithelial (*CDH1*) [12], mesenchymal *snai1* and epithelial *sox10* in zebrafish [13], or mesenchymal *Vimentin (Vim)* in mice [14]. Nevertheless, heterologous expression of exogenous reporter genes is hampered by the existence of alternative promoters, *cis*- and *trans*-regulatory elements, and epigenetic events that modulates promoter activation making some data unreliable.

To overcome these limitations, we genetically engineered and characterized human lung carcinoma H2170 Vimentin Reporter Cells (VRCs) where the fluorescent protein coding gene (*mCardinal*) has been knocked-in as a genetic fusion, although separate proteins due to a T2A self-cleaving peptide, at both alleles of the mesenchymal marker *VIM*.

## 2. Materials and Methods

### 2.1 Cell culture, nucleofection and transfection

Human squamous lung carcinoma (H2170), human colorectal adenocarcinoma (HT29) as well as human embryonic kidney (HEK293) cells were cultured according to standard mammalian tissue culture protocols and sterile techniques. The cell lines were cultured in DMEM supplemented with 10% fetal calf serum (FCS), 100 units/mL and penicillin/100 µg/mL streptomycin. All tissue culture media and supplements were obtained from Gibco. H2170, HT29 and HEK293 cells were obtained from ATCC. H2170 cells were nucleofected for genome editing with the use of Nucleofector I Device and Cell Line Nucleofector Kit T (Amaxa). The optimized protocol is as follows: nulceofection of 1 million suspended cell with 2 µg of the plasmid DNA using program X-001, generating transfection efficiency of 99%. For performing functional experiments on a smaller scale the H2170 cells were transfected with the use of Lipofectamine 3000 (Invitrogen) with transfection efficiency 90-95% after 24 h. 50,000 cells were seeded a day before transfection in 24-well plates. Transfection was performed using 500ng of the plasmid DNA, 1.5 µL Lipofectamine®3000 Reagent and 1 µL P3000™ Reagent (both from Invitrogen) per well, following the manufacture’s protocol. HEK293 were transfected with the use of TurboFect reagent (Thermo) according to the protocol supplied by the company. Next day, the transfection efficiency was in range 80-90%.

### 2.2 Plasmid vectors for the knock-in to VIM

We modified Cas9/gRNA expressing plasmid pSpCas9(BB)-2A-Puro (PX459) V2.0 (Addgene plasmid #62988) by inserting the fragment coding the targeting gRNA using digestion/ligation protocol [15]. The oligonucleotides used to generate the gRNA were gRNA-F and gRNA-R (Table S1). The template plasmid used for inserting DYKDDDDK-tagged (FLAG) mCardinal fluorescent protein gene after *VIM* was designed as following (gRNAsite-800nt of *Vim*-P2A-mCardinal-FLAG-800nt of 3’*Vim*UTR-gRNAsite) and was synthetized by Thermo. The Cas9-gRNA and the template plasmids were both nucleofected to cells at the same time and the cells were selected 2 days upon nucleofection for another 2 days in 5 µg/ml of puromycin. The positive single cell clones obtained by dilution were genotyped what was described in details in Genotyping section.

### 2.3 Plasmid vectors used in functional assays

For functional experiments in H2170 knocked-in cells we used (FLAG) Snail 6SA (active Snail) plasmid which was a gift from Mien-Chie Hung (Addgene plasmid # 16221) [16], TGFB1-bio-His (proTGFβ) which was a gift from Gavin Wright (Addgene plasmid # 52185) [17] and HA-OVOL2 (OVOL2) expressing plasmid was a kindly gift from Changwon Park [18]. EMT/MET in VRCs was studied with the use of expressing vectors harbouring genomic fragments of microRNA-200 family (miR-145, miR-200b, miR-200c, miR-205) which were cloned in our laboratory (See Molecular Cloning section). Control cells were transfected with pUC18 sub-cloning plasmid.

### 2.4 Molecular cloning

All the plasmid fragments used for cloning were amplified using tiHybrid proofreading DNA polymerase (EURx), according to the supplied protocol. PCR products amplified on the plasmid DNA template were incubated overnight at 37 °C with DpnI FastDigest enzyme (Thermo) as in manufacturer’s instructions.

*VIM-T2A-mCardinal* sequence was cloned from cDNA of knocked-in cells upon PCR amplification into linearized by PCR pmR expressing vector (Clonetech) and recombined using Gibson Assembly reagent (NEB). Resulting vector contained full *VIM-T2A-mCardinal* reading frame under the control of CMV promoter. The primers for insert amplification were KI-F and KI-R whereas the pair used for backbone linearization were BCB-F and BCB-R (Supplementary data, Table 1). Mutagenesis was performed by REPLACR methodology [19], using the SDM-F and SDM-R primers (Suppl material Table 1).

The vectors harbouring miR-145, miR-200b, miR-200c and miR-205 genomic fragments were created by inserting each PCR amplified microRNAs gene into the 3’UTR of mNeon fluorescent protein expressing vector (pmR-mNeon). All listed above genomic fragments were amplified using tiHybrid DNA polymerase (EURx) from the DNA, which was purified from the blood of healthy volunteer with the use of GeneAll Exgene Blood SV kit (GeneAll). The sets of primers used for amplification of miR-145, miR-200b, miR-200c and miR-205 fragments were miR145-F/miR145-R, miR-200b-F/miR-200b-R, miR-200c-F/ miR-200c-R, miR-205-F/ miR-205-R respectively (Suppl material Table 1.). The amplified products produced proper sticky ends upon digestion by BglII and HindIII restrictases (both from Thermo). Digested and purified DNA fragments were ligated using T7 ligase (Thermo) in molar ratio 3:1 with 100 ng of linear pmR-mNeon, which was previously cut by BglII and HindIII enzymes.

Resulting vectors were named miR-145, miR-200b, miR-200c, miR-205. The sequences of all the vectors were verified by Sanger sequencing (Genomed).

### 2.5 Genotyping

The targeting sequence for CRISPR/Cas9 was in the last intron (intron 8) of *VIM* in HEK293 and H2170 cells with the use of Benchling algorithm. Single-cell clones were cultured on 96-well plates to obtain more than 50% of confluence (Nunc). Upon washing with Phosphate-buffered saline (PBS, Gibco) they were genotyped by PCR using Mouse Direct PCR Kit (Bimake), following the manufacturers instruction. The primers used for genotyping were 170 and 249 (Supplementary data, Table 1). The genotyping was further confirmed by PCR using the same set of primers (170 and 249), tiHybrid DNA polymerase and high quality genomic DNA, purified from single-cell clones with the use of QIAamp DNA Mini Kit (Qiagen). Then the KI was verified by sequencing (Genomed).

### 2.6 RNA extraction, reverse transcription and qPCR

Total RNA was isolated from cells with the use of Extractme Total RNA kit (Blirt) according to manufacturer’s manual, including DNase treatment. The purity and quantity of isolated RNA was estimated spectrophotometrically with the use of Tecan M200Pro microplate reader supplied with NanoQuant plate (Tecan). Only the samples with 260/280 nm OD ratio higher than 1.8 were used for downstream analysis. For molecular cloning 3 µg of RNA were reverse transcribed for 30 min at 50°C using an oligo(dT) primer and the Transcriptor High Fidelity cDNA Synthesis Kit (Roche) followed by 5 min enzyme inactivation at 85°C according to manufacturer’s instructions. For QPCR, 2 µg of RNA were reverse transcribed using High-Capacity RNA-to-cDNA Kit (Applied Biosystems) following to the manufacturer’s protocol.

Quantitative real-time expression analysis was performed using a LightCycler®480 II instrument (Roche) equipped with 384 well plates and PowerUp™ SYBR™ Green Master Mix (Applied Biosystems). The primers were as listed in Supplementary data (Table S1). Amplification was performed in 12,5 µl reaction mixture containing cDNA amount corresponding to 12.5 ng of total RNA, 1 x PowerUp SYBR Green Master Mix and 3.125 pmol of each primer (Forward and Reverse). Upon 2 min of initial incubation at 50 °C followed by 2 min incubation at 95°C, cDNA was amplified in 45 cycles consisting of 15 s denaturation at 95°C, 30 s annealing at 60°C and 20 s elongation at 72°C. Obtained fluorescence data was analyzed using a relative quantification (RQ) method 2^(–ΔΔCT) for estimating expression fold changes normalized to dim-VRCs and 2^(–ΔCT) method for comparison of the expression of each measured gene. The assessed genes expression (*VIM, mCardinal*, C*DH1, ZEB1* and *ZEB2)* were normalized to *GAPDH* level, which was measured with the use of GAPDH-F and GAPDH-R oligonucleotides. *GAPDH* was previously confirmed as stably expressed at mRNA level in H2170 as well as in VRCs (mean Cp = 17.14; median Cp = 17.09; SD = 0.5209; SEM = 0.03638; N = 205). Expression of *VIM* was measured using VIM-F and VIM-R primers, measurement of *mCardinal* level was conducted using mCard-F and mCard-R primers, whereas estimation of *CDH1* expression was performed at mRNA level with the use of Cdh1-F and Cdh1-R oligonucleotides. *ZEB1* and *ZEB2* quantifications were performed using ZEB1F/R and ZEB2F/R pairs of oligonucleotides, respectively. *TWIST1* and *TWIST2* quantifications were performed using TWIST1F/R and TWIST2F/R pairs of oligonucleotides, respectively. The primers were designed as intron-spanning to avoid any influence of genomic DNA contamination and was listed in Supplementary materials (Table S1).

### 2.7 Flow cytometry of living cells

VRCs or HT29 cells cultured for one day on 6 well plate (Nunc) were washed with HBSS - Hank’s Balanced Salt Solution (Thermo) and detached by Accutase (Corning). The cells dissolved in 100 µl HBSS with 5 µl of PE Mouse anti-E-Cadherin (562526, BD Pharmingen) were incubated for 1 hour, washed with HBSS and suspended in 0.5 ml of HBSS. The cells were counted on FL-3 fluorescence channel using FACSCalibur flow cytometer (BD).

### 2.8 Cell sorting

VRCs seeded one day before were then sorted using the BD FACSAria™ flow cytometer (BD Biosciences). In this case, cells were selected into two populations due to the intensity of their mCardinal fluorescence in farred channel (Figure S1). After sorting, the purity of sorted cells was confirmed by flow cytometry and reaches more than 97%. The resulting two populations of VRCs were named dim-VRCs and bright-VRCs.

### 2.9 Confocal imaging of living cells

The cells were seeded onto 24-well glass-bottom plates (MoBiTec) a day before transfection. Transfection was conducted as described above. Transfected cells were then visualized under Nikon Ti Confocal microscope in 24 and 28h upon transfection using 563nm laser for mCardinal fluorescence. Mean fluorescence intensity was measured using NISelements (ver 3.22.08) software and shown on the graphs. Each of at least 10 measured Regions of the interest (ROIs) included the cluster of more than 20 adherent cells. The brightest slice of the z-stack were chosen for the measurement. The measurement were done in duplicate.

### 2.10 Immunocytochemistry

The cells seeded a day before on glass bottom Labtec chamber slides (Nunc) were washed with PBS and fixed with 2% paraformaldehyde for 20 minutes in room temperature. Upon washing in PBS the cells were permeabilized by 0.5% triton in PBS for 20 min. The endogenous peroxidase was blocked by incubation in 1% sodium azide and 1% hydrogen peroxide in PBS for another 20 min. Another washing step in PBS was followed by incubation in Blocking Buffer which was 1x Normal Donkey Serum (reconstituted from 20x, Jackson ImmunoResearch) containing 1% bovine serum albumin (SantaCruz Biotechnology) and 0.1% of triton for 30 min in room temperature. Supplied by Cell Signaling primary antibodies against VIM (rabbit, #5714S) and FLAG-tag (mouse, #8146S) were diluted in Blocking Buffer in ratio 1:100 and 1:1600 respectively. Incubation with the primary antibodies was conducted overnight at 4°C and was followed by washing in PBS. Secondary Peroxidase F(ab’)_2_ Fragment Donkey Anti-Rabbit IgG (H+L) (711-036-152, Jackson ImmunoResearch) for detection of primary rabbit and Peroxidase F(ab’)_2_ Fragment Donkey Anti-Mouse IgG (H+L) (715-036-150, Jackson ImmunoResearch) for detection of primary mouse antibodies were diluted in Blocking Buffer 1:1000 respectively. Incubation with secondary antibodies (fluorochrome or peroxidase conjugated) was occurred for 1 h in room temperature and was followed by another washing in PBS. Peroxidase conjugated antibody were stained with Alexa Fluor 555 Tyramide Reagent (B40955, LifeTechnolgies) or Alexa Fluor 488 Tyramide Reagent (B40953, LifeTechnolgies) following to the manufacturers protocol, resulting in enzyme-linked fluorescence signal amplification. When double Alexa Fluor Tyramide Reagents (488 and 555) staining was done an additional step, 15 minutes washing in 3% hydrogen peroxide, 0.1% sodium azide in PBS, to achieve complete inhibiting of HRP enzyme was added between incubations with two different secondary HRP-antibodies. The cells were finally washed by PBS and stained with Hoechst (Cayman) diluted in PBS (1:1000 from 10 mg/mL stock solution). The cells were washed once again, mounted with ProLong Gold mounting medium (LifeTechnologies) and visualized under Nikon Ti confocal microscope.

### 2.11 Migration of VRCs

XCELLigence real-time cell analysis system (Acea) equipped with 16-wells CIM-plates served for VRCs migration assessment. VRCs cells first detached using Accutase, then transfected with OVOL2, miR-200c, miR-205 or pUC18 plasmids using Lipofectamine300 reagent, as was described above. Then, 30,000 of cells were seeded on each well of the CIM-plate, kept in the standard mammalian tissue culture conditions and measured for 24 hours. The cell migration toward an attractant, which was 10% DMEM supplemented with 10% FCS was normalized to cell migration toward DMEM alone and shown as Cell Index. Each experiment was measured at latest in triplicates and repeated twice.

## 3. Results

### 3.1.1 Genome editing

The targeting sequence in H2170 cells was confirmed by sequencing (Figure S1). The H2170 lung cancer cells were nucleofected with the CRISPR and template (vKIT, Figure 1A) plasmids, following enrichment using puromycin. vKIT contained the sequence of self-cleaving T2A peptide followed by sequence of FLAG-tagged *mCardinal* fluorescent protein gene, flanked by two 800 nucleotides long homology arms (HA) that corresponds to *VIM* gene fragments near CRISPR-targeting region (Figure S2).

**Figure 1.**
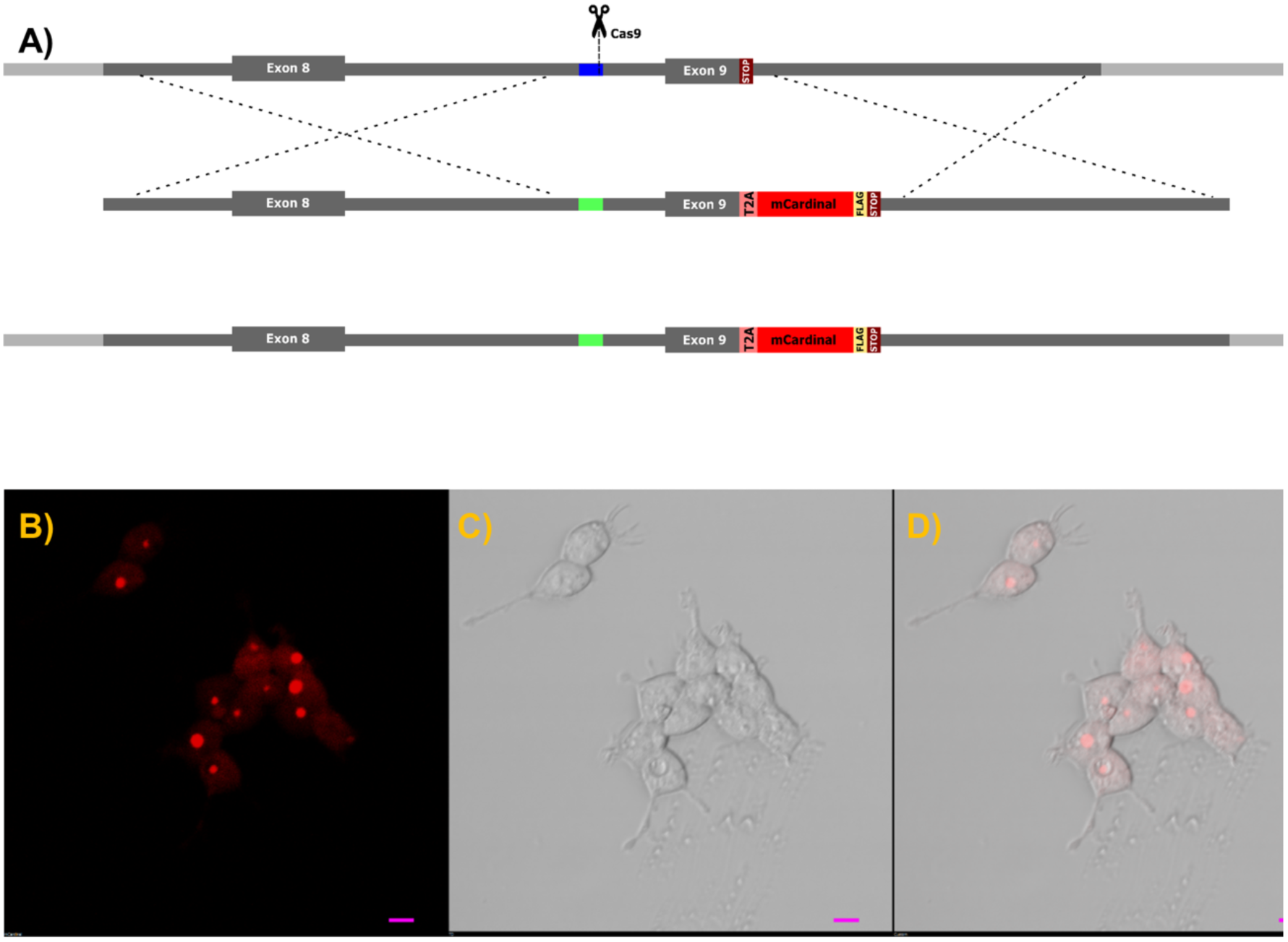
**A)** Schematic representation of genome editing strategy. Genomic region of H2170 cells targeted by CRISPR/Cas9 with homology arms marked in grey while targeted site in intron 8 has been marked in dark blue. The donor DNA template contains the sequence of T2A-mCardinal-FLAG, flanked by two homology arms that corresponds to VIM gene fragments (grey) recombines with targeted gene. VIM allele in VRCs contains the knocked-in DNA and changed sequence recognized by Cas9 sequence is coloured in green. **B-D)**. Subcellular localization of mCardinal fluorescent protein in living VRCs. The scale bar is 10 µm. **B)** mCardinal, **C)** transmission, **D)** merged.

The template cassette, containing *HA-T2A-FLAG-mCardinal-HA*, was flanked by two gRNA targeting sequences identical to those for *VIM* gene in order to linearize the knock-in insert and promote efficient Homology directed repair (HDR).

Two days after nucleofection, the cells were selected using puromycin (1 µg/ml) for another two days. Single cell clones were obtained by dilution, further they were genotyped by PCR with the efficiency near 2,7% (3/112). Single cell clones, confirmed by PCR, were called VRCs and used in downstream analysis. The sequence of knocked-in DNA fragment in the VRCs was further verified by Sanger sequencing (Figure S1).

The subcellular localisation of mCardinal in living cells (Figure 1B-D) by confocal imaging showed that mCardinal was present in the cytoplasm, but its concentration was not homogenous showing bright foci as well as cytoplasmic localisation. Immunofluorescent labelling of VIM and FLAG-mCardinal (Figure 2A-F) shows almost complete co-localization, where VIM was found in the cytoskeleton as well as in the foci.

**Figure 2.**
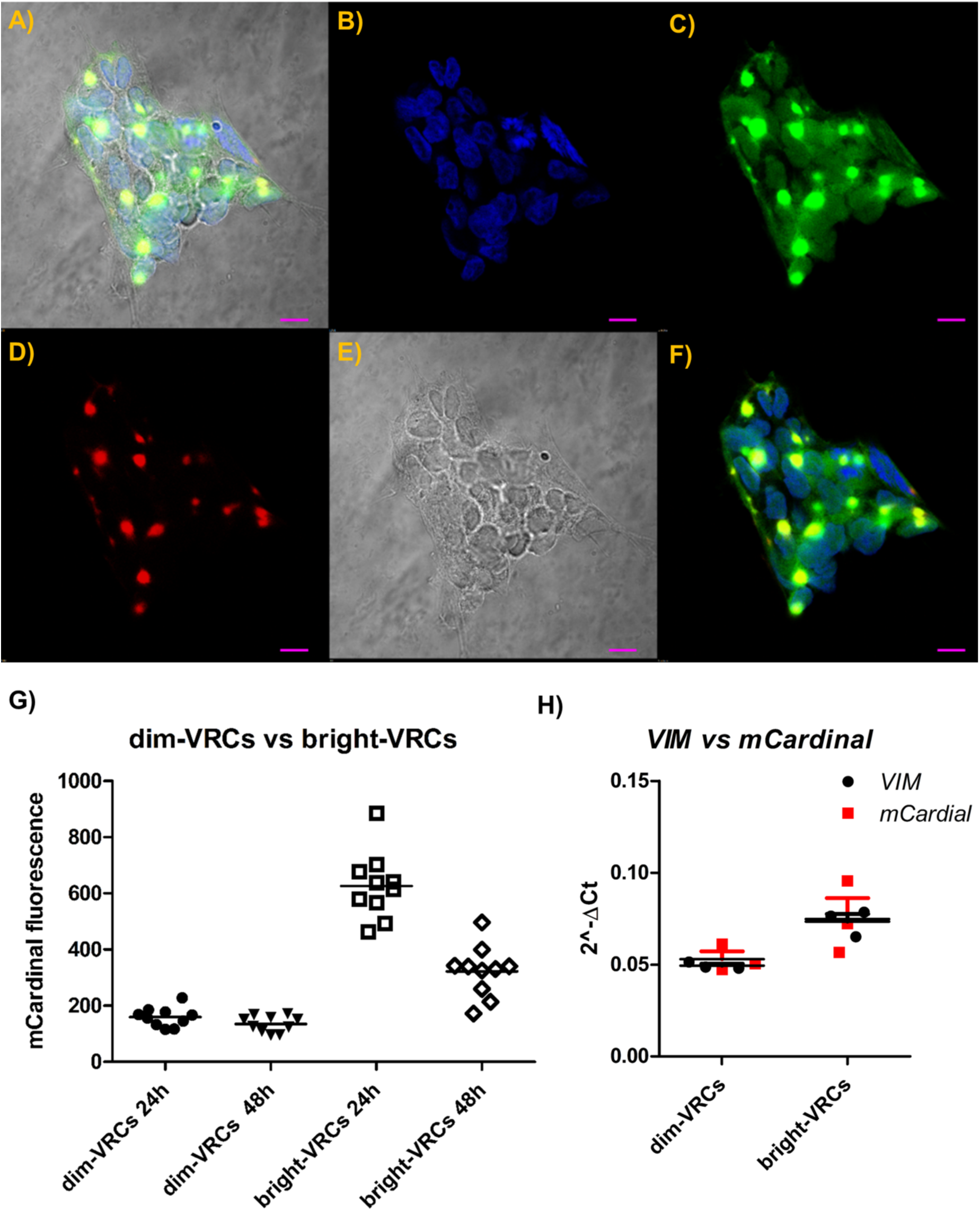
**A-F)** Immunofluorescent labeling of FLAG-tagged mCardinal and VIM in VRCs. Shown in green FLAG tag **(C)** is seen as bright spot-like areas surrounded by less intensive fields localized in the cytoplasm. Localization of VIM shown in red **(D)** corresponds to the regions with the highest fluorescence of FLAG-tagged mCardinal **(C)**. Blue coloured DAPI, FLAG, VIM and transmission were merged on picture **A). B)** DAPI, **C)** anti-FLAG, **D)** VIM, **E)** transmission, **F)** merged fluorescence. The scale bar is 10 µm. G) mCardinal fluorescence changes of sorted VRCs populations with time. The fluorescence of dim-VRCs and bright-VRCs populations was measured in 24 h and 48 h of culture upon sorting. The points correspond to the fluorescence of selected ROI, whereas the lines show mean fluorescence. The representative results was shown. **H)** *VIM* and *mCardinal* quantification by qPCR shows similar amount of both transcripts. VIM and mCardinal genes were measured in dim-VRCs transfected cells in 24 or 48 h upon transfection. The data shows mean 2^(–ΔCT) relative to *GAPDH* housekeeping gene. The graph shows the representative result of the measurement which was done in triplicate.

The unexpected cellular localization of VIM made us to analyze if the gene product was intact. For this we amplified the entire *VIM-T2A-mCardinal* from mRNA, and cloned it into an expression vector. The whole *VIM-T2A-mCardinal* sequence was sequence-verified. This VIM-T2A-mCardinal cDNA exerted identical cellular distribution of mCardinal under the confocal microscope when expressed in HEK293 cells (Figure S3 A-C).

To determine if fusing mCardinal to VIM would result in a better cellular distribution, the T2A peptide sequence was modified by substitution of the P16A and P18A in the T2A sequence. The resulting vector (*VIM-mCardinal*) was overexpressed in HEK293 cells, showing granular localisation too (Fig S3 D-F). Overexpression of VIM-mCardinal in HEK293 cells showed that this fusion protein localised only in large granules in the cell and is similar in comparison to the fluorescence in VIM-T2A-mCardinal overexpressing cells, where the fluorescence was shown in similar granules as well as diffusing in the cytoplasm (Figure S3).

Since the foci in the Vim-T2A-mCardinal cells corresponded to fused Vim-mCardinal fusions, likely resulting from inefficient T2A processing, were brighter than if evenly distributed throughout the cytoplasm, we considered them as a baseline.

To further characterise VRCs, they were then sorted according to the fluorescence intensity, resulting in two populations (dim-VRCs and bright-VRCs) that differ about 3 times in mCardinal fluorescence intensity (Figure S4). That difference was reduced upon culturing to less than 2 fold mCardinal intensity between dim-VRCs and bright-VRCs after 48 h (Figure 2G). *VIM* as well as *mCardinal* expression measured at the transcriptional level confirmed that those two genes are expressed at equal levels (Figure 2H and S5).

The dim-VRCs where then exposed to a series of EMT-inducing factors, to analyse the correlation between mesenchymal conversion and mCardinal expression.

### 3.1.2. EMT markers vs reporter gene

The dim-VRCs were transfected with active *Snai1* or proTGFβ expressing plasmids, which resulted in a 2-fold increase in fluorescence, examined using confocal microscopy in 24 h or in 48 h after transfection (Figure 3A). Correspondingly, the transcriptional levels of *VIM* as well as *mCardinal* showed similar fold increase (RQ) (Figure 3B-C). The highest expression of *VIM* was seen in the cells that overexpressed active Snai1 one and two days upon transfection (median RQ = 1.625 and RQ = 1.580 respectively). proTGFβ overexpressing dim-VRCs for 24 h and 48 h also overexpressed *VIM* at median level RQ = 1.371 and RQ = 1.424. Control population of bright-VRCs expressed *VIM* at median RQ = 1.569. The highest level of *mCardinal* was expressed in active Snai1 expressed dim-VRCs after 24 h (median RQ = 1.357) and in 48 h after transfection (median RQ = 1.240). proTGFβ overexpressing dim-VRCs for 24 h and 48 h also overexpressed mCardinal at median level RQ = 1.206 and RQ = 1.248, whereas bright-VRCs *mCardinal* median RQ was 1.181. The expression levels of *VIM* as well as *mCardinal* relative to *GAPDH* housekeeping gene show a similar pattern (Figure 3B-C). Epithelial marker *CDH1* transcript in proTGFβ over-expressing dim-VRCs did not reveal significant changes in 24 h nor in 48 h upon transfection (Figure 3D). We did not observed significant changes *CDH1, VIM, SNAI1, ZEB1, ZEB2, TWIST1* or *TWIST2* level between VRCs and parental H2170 cells by qPCR (Figure S6).

**Figure 3.**
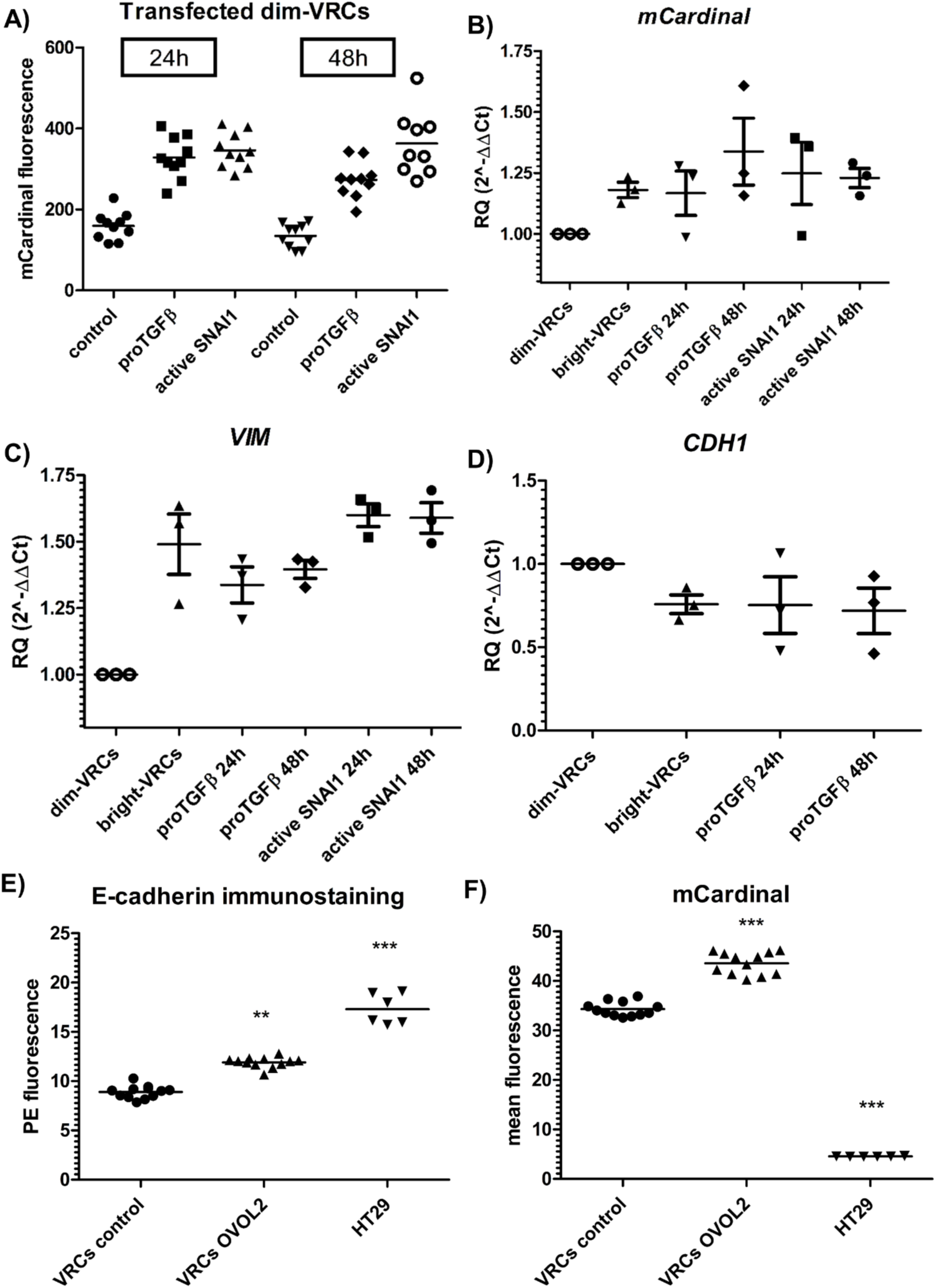
A) mCardinal fluorescence of dim-VRCs population transfected with active SNAI1 or proTGFβ plasmids. The fluorescence of the cells was measured in 24 h and 48 h of culture upon sorting. The points correspond to the fluorescence of selected ROI, whereas the lines show mean fluorescence. The representative results was shown. **A)** *mCardinal* and B) *VIM* relative quantification (RQ) by qPCR. VIM and mCardinal genes were measured in dim-VRCs transfected cells in 24 or 48 h upon transfection. The data shows mean 2^(–ΔΔCT) relative to *GAPDH* housekeeping gene. The results were normalized to control which were pUC18 transfected dim-VRCs cells. The graph shows the representative result of the measurement which was done in triplicate. **D)** Relative quantification of *CDH1* by qPCR. *CDH1* levels were measured in proTGFβ transfected dim-VRCs in 24 or 48 h upon transfection. The graph shows mean RQ 2^(–ΔΔCT) ± SEM relative to *GAPDH* housekeeping gene. The results were normalized to control which were pUC18 transfected dim-VRCs cells 48 h upon transfection. The graph shows the representative result of the measurement which was done in triplicate. No effect in CDH1 expression. **E)** Immunostaining of E-cadherin and F) mCardinal fluorescence in VRCs and HT29 E-cadherin positive cells by flow cytometry. The graph shows: VRCs control were VRCs transfected with pUC18 vector whereas, VRCs OVOL2 were OVOL2 overexpressing in 24 h upon transfection. Immune-stained un-transfected HT29 cells were used as controls. The graph shows the single reads as well as the mean value of 3 independent experiments. Statistical significance in comparison to control was calculated using Mann-Whitney test and was rated by asterisk: * p>0.05; ** p 0.01; *** p<0.01.

### 3.1.3. OVOL2 and miRNAs overexpression modulates CDH1 expression in VRCs via ZEB1/2 repressors

In order to test if mesenchymal-like VRCs could reverse their phenotype into more epithelial, we transfected these cells with a strong epithelial activator - OVOL2 [20]. Immunostaining followed by flow cytometry analysis as well as qPCR confirmed our hypothesis showing that E-cadherin level is elevated in OVOL2 overexpressing VRCs after 48h (Figure 3E). OVOL2 overexpressing VRCs concomitantly showed a significant increase of the mCardinal fluorescence (Figure 3F). To examine the mechanism of epithelial phenotype development by VRCs, we analysed the expression level of CDH1, as well as its negative regulators *ZEB1* and *ZEB2* (Figure 4A-C) in the cells that overexpressed OVOL2 or four different *ZEB1/2*-targeting members of the miR-200 family (miR-145, miR-200b, miR-200c, miR-205). Both negative regulators of *CDH1* expression (*ZEB1* and *ZEB2*) were significantly downregulated in VRCs that overexpressed OVOL2 or miR-205 with simultaneous upregulation of *CDH1*. miR-200c overexpression in VRCs produced increased level of *CDH1* together with downregulated ZEB1, whereas the effect of miR-145 overexpression was seen only as elevated *CDH1* level. We did not observe changes in *CDH1* or *ZEB1/2* levels in mirR-200b-expressing VRCs.

**Figure 4.**
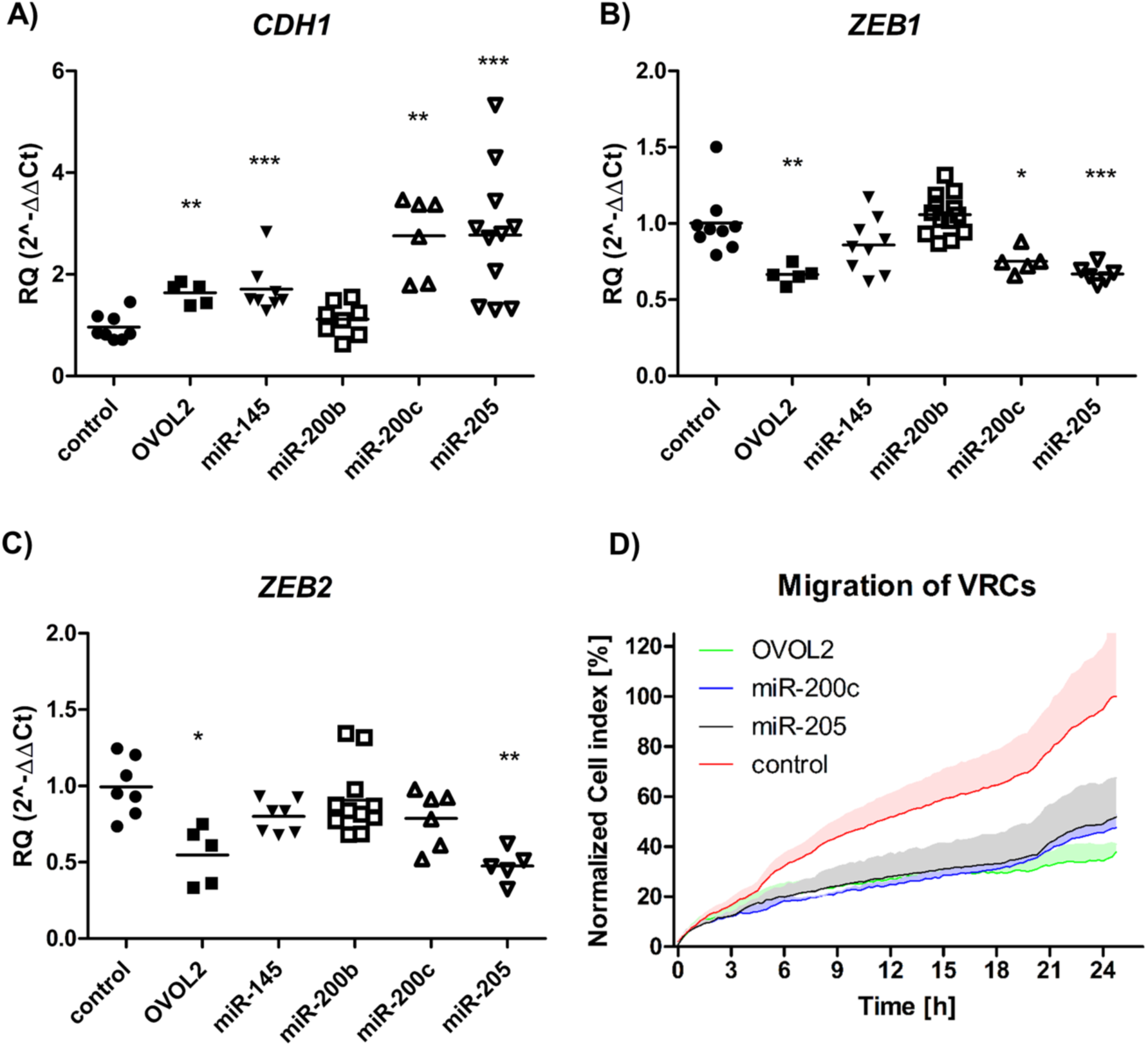
**A-C)** Relative quantification of *CDH1, ZEB1* and *ZEB2* transcripts in transfected VRCs by qPCR. The expression of genes relative to *GAPDH* housekeeping gene was measured in 48 h upon transfection of VRCs with *OVOL2* and microRNA expressing vectors (*miR-145, miR-200b, miR-200c* and *miR-205*). The results were normalized to VRCs control (mock transfected) and shown as mean RQ 2^(–ΔΔCT) as well as single values. The graph contains data from at least two independent experiments which were measured in triplicate. Statistical significance in comparison to control was calculated using Mann-Whitney test and was rated by asterisk: * p>0.05; ** p 0.01; *** p<0.01. **D)** Real-time migration analysis of transfected VRCs using xCELLigence system. The line shows mean normalized cell index, whereas coloured area visualizes standard deviation of 3 replicates. Migration of VRCs transfected with: *OVOL2* is shown in green, *miR-200c* shown in blue, *miR-205* shown in black, whereas pUC18 as the control is coloured in red. The representative results from two different experiments was presented.

### 3.1.4. OVOL2 and miR-200c or miR-205 overexpressing VRCs show decreased migratory potential

In order to examined if decreased levels of ZEB1/2 could repress migratory properties of VRCs [20,21], the cells were transfected with *OVOL2, miR-200c* and *miR-205* overexpressing plasmids and real time migration assay was conducted. We observed over two fold decrease in the number of migrating VRCs that overexpressed all of tested factors that may modulate ZEB1/2 (OVOL2, miR-200c and miR-205) in comparison to the mock transfected cells (Figure 4D).

*3.2. Figures*

## 4. Discussion

Mounting evidence shows that tumors are far more heterogenous than expected in due to genetic diversity of the tumor cells as well as their phenotypic plasticity [22,23]. This phenotypic plasticity is in fact a phenotype switching, what is understood as a phenomenon whereby cancer cells transit between different phenotypes in response to environmental cues, without acquiring new mutations [24]. This cellular plasticity has been reported as having high clinical impact because it is crucial for drug resistance development e.g. in lung cancer patients treated with EGFR inhibitors [25], re-acquiring pluripotency and become a cancer stem cell (CSC) [26], or maintaining metastatic ability of many types of cancer cells [27–29].

Among tumor cells there are some that undergo EMT, MET, or hybE/M, and in order to understand the effects of cellular plasticity in biological and pathological processes, there is a need of reporters that can determine the stage of a cell. Observing changes in mesenchymal and epithelial phenotype as they occur, is a significant improvement in studying of molecular mechanism in cancer cells. H2170 cells was chosen because they have been a good model for to EMT and MET [30,31].

To-date EMT/MET is routinely studied by the use of exogenous reporter genes that are heterologously expressed exhibiting interference of cis- and trans-regulatory elements, alternative promoters or epigenetic events that resulted in obtaining unreliable data [32,33]. These limitations are overcome by using genome engineering of endogenous genes. There are a couple of available cancer cell lines that harbor a C-terminal red fluorescent protein (RFP) tag on VIM: A549 (lung), HCT116 (colorectal), MDA-MB-231 (breast adenocarcinoma) and they are commercially available at ATCC cell repository. These cells however, have the *RFP* knocked-in into the beginning of last exon of *VIM* (exon 9), what results in deletion of large fragment of that exon and of the gene product (https://www.lgcstandards-atcc.org/en/Global/Products/CCL-247EMT.aspx#documentation). While, these modified A549, HCT116 or MDA-MB-231 cell lines enable near real-time tracking of the EMT/MET status as cells transition from epithelial to mesenchymal phenotype under defined conditions [34], any approach where the an endogenous protein is truncated may affect functions in comparison to endogenously expressed proteins,, in particular in the context of a protein such as VIM which has many signaling modulatory activities during cell plasticity [35–37].

Using a far-red fluorescent protein (mCardinal) which is simultaneously expressed with endogenous VIM, but separate during translation due to the viral self-cleaving peptide (T2A). T2A was chosen because, together with P2A, it have been reported as the most efficient from all tested self-cleaving 2A peptides [38–41], moreover T2A resulted in the least amount of “uncleaved” protein product among the family of 2A peptides [39–41]. The stability and half-life of the 2 resulting proteins can result in small changes in total expression, the two proteins are synthesized at a 1:1 ratio [41]. *VIM* expression at the transcriptional level corresponded to that of *mCardinal*, and when stimulated, the changes of *VIM* expression when hand-to-hand to those of *mCardinal*, evidencing that our strategy is fully functional.

mCardinal was chosen by us mainly because this far-red monomeric protein is better for *in vivo* settings due to it is excitation as wavelengths above 600 nm, which penetrate through hemoglobin-rich tissues far better than lower wavelengths [42]. The second reason is that the mCardinal is suitable for precise monitoring of its changing expression in due to its short maturation half-time (27 min) [43] which is about 3 times shorter when we compare others commonly used fluorescent markers e.g. RFP [44].

The subcellular localisation of mCardinal in VRCs by confocal imaging show non-homogenous distribution of the fluorescence which was observed at moderate level in cytoplasm as well and in brighter glowing foci. Immunofluorescent staining of VIM and FLAG-tag mCardinal showed co-localization at the “foci”, whereas Vimentin is also present in the cytoskeleton of VRCs. Moreover, re-cloning of and the *VIM-P2A-mCardinal* and further over-expressing it in HEK293 cells shown similar sub-cellular distribution of the fluorescence as in VRCs. The presence of foci is probably the result of not fully efficient T2A peptide is cleavage, the fusion protein being unable to form normal VIM polymers [39–41], which is more conspicuous in the case of the fusion of VIM-mCardinal. In fact, C-tags in VIM might result in an impaired protein [45], and it’s expression may give dominant-negative phenotype, similar to the phenotype observed in the cells which express only the N-terminal domain of VIM (NTD-VIM) [46]. NTD-VIM domain (VIM that lacks C-terminus) expressing cells show its abnormal localization, as formation of the granules in the cytoplasm together with disrupted endogenous VIM localization examined by immunostaining [45]. The role of the tail domain in VIM organisation is not fully understood, although it is known to be necessary for appropriate network formation [47]. Thus, it seems to be possible that abnormal VIM localisation in VRCs may be produced by the fluorescent protein. Yet, the presence of bright foci is advantageous as these structures are brighter than diffused cytoplasmic mCardinal under low VIM expression.

Dim-VRCs presented a reduced mesenchymal phenotype as compared to bright-VRCs. The dim-VRCs could be driven to a more mesenchymal phenotype via expression of proTGFβ [17] or constitutively active SNAI1 transcription factor [16]. We also noticed insignificant downregulation of *CDH1* in proTGFβ over-expressing dim-VRCs, what suggests that E-cadherin may be only partially regulated via TGFβ-dependent pathway [48].

*OVOL2* overexpressing VRCs respond by E-cadherin expression, what is in accordance with data describing OVOL2 as a strong epithelial regulator that maintains transcriptional programs in epidermal keratinocytes and mammary epithelial cells by repressing ZEB1 and ZEB2 [49,50].

MiR-200 family were identified as double-negative regulators by silencing *CDH1* and transcription repressors Zeb1 and Zeb2 [51,52]. We confirmed the activatory role of miR-145, miR-200c, miR-205 on *CDH1* expression in VRCs. We also confirmed that miR-200c expression decreased *ZEB1* level in VRCs [53], but we did not observe any effect on miR-200c on *ZEB2* expression [54]. VRCs overexpressed miR-205 shown *ZEB1/2* downregulation in cancer cells, what is in accordance with previous works [55,56]. Our results, suggests that the molecular mechanisms of miR-200 family on EMT might be cell type specific and more work needs to be done to clarify this. Our data is in accordance with previous data in that the migratory potential of mesenchymal cells is inhibited by ZEB1 or ZEB2 downregulation produced by miR-200 family overexpression [57].

The very similar level of *CDH1, VIM, SNAI1, ZEB1, ZEB2, TWIST1* or *TWIST2* transcript between VRCs and parental H2170 cell line suggests that VRCs may be considered as the model that is useful in broad range of studies.

In conclusion, our data uniquely illustrate a reporter line, VRCs, as a reliable reporter model for studying EMT and MET. VRCs allow to directly observe of cellular plasticity with respect to their mesenchymal/epithelial *in vitro* and in the future *in vivo*. These cells could also be used as a robust platform for drug development.

## Author Contributions

Conceptualization, ARM, JCz; Methodology, MK, JCz, ARM; Validation, MK, JW; Formal Analysis, MK; Investigation, MK; Resources, AS, ARM; Writing – Original Draft Preparation, MK; Writing – Review & Editing, MK, JCz, JK; Visualization, MK; Supervision, ARM, AS; Project Administration, ARM; Funding Acquisition, ARM.

## Acknowledgments

We would like to thank to prof. Jacek Roliński for enable of the use of his lab equipment.

