## Supplementary data for "Genetical engineered lung cancer cell for analyzing Epithelial-Mesenchymal transition"

**Figure S1.** Alignment of the WT, expected and actual sequences encoding Vim (NM_003380) and the knock-in region. The expected KI and actual sequences matched perfectly while they differ to the WT in the expected regions. Two reads amplified using each of the following primers: 170F, 324R, 324F and 170R had been shown. Recoded bases corresponding to gRNA targeting site were coloured in red.

5'HA; VIM missing part; T2A; mCardinal; 3xFLAG; 3'HA; VIM gRNA target site

```
expected    TCGAACCCAAGTACACTCTTGCATTCTATGCTTTAAGTTAAATGCAAACTCCTTTTTCCT 60
genomic     --------------------------------------TAAATGCAAACTCCTTTTTCCT 60
170FI       ------------------------------------------------------------ 60
170FII      ------------------------------------------------------------ 60
324RI       ------------------------------------------------------------ 60
324RII      ------------------------------------------------------------ 60
324FI       ------------------------------------------------------------ 60
324FII      ------------------------------------------------------------ 60
170RI       ------------------------------------------------------------ 60
170RII      ------------------------------------------------------------ 60

expected    TCTTCCTGCTGCAAGTACTATCTCATCCTGATGCTCAAGAGTGTCAGGGCCTGGGTTTCC 120
genomic     TCTTCCTGCTGCAAGTACTATCTCATCCTGATGCTCAAGAGTGTCAGGGCCTGGGTTTCC 120
170FI       --------------------TCTCATCCTGATGCTCAAGAGTGTCAGGGCCTGGGTTTCC 120
170FII      ---------------------CTCATCCTGATGCTCAAGAGTGTCAGGGCCTGGGTTTCC 120
324RI       ------------------------------------------------------------ 120
324RII      ------------------------------------------------------------ 120
324FI       ------------------------------------------------------------ 120
324FII      ------------------------------------------------------------ 120
170RI       ------------------------------------------------------------ 120
170RII      ------------------------------------------------------------ 120

expected    AAACAGAGACTACCCTAAAATTATTTGGCGAGTAGTACTTTACACAATTGCCTCTCCCCC 180
genomic     AAACAGAGACTACCCTAAAATTATTTGGCGAGTAGTACTTTACACAATTGCCTCTCCCCC 180
170FI       AAACAGAGACTACCCTAAAATTATTTGGGGAGTAGTACTTTACACAATTGCCTCTCCCCC 180
170FII      AAACAGAGACTACCCTAAAATTATTTGGGGAGTAGTACTTTACACAATTGCCTCTCCCCC 180
324RI       ------------------------------------------------------------ 180
324RII      ------------------------------------------------------------ 180
324FI       ------------------------------------------------------------ 180
324FII      ------------------------------------------------------------ 180
170RI       ------------------------------------------------------------ 180
170RII      ------------------------------------------------------------ 180

expected    ACAAATCATAATTGTTTCAGTAAAATGGTTACTTGGTTTTTCCAAGAAAAAACTCGTTTT 240
genomic     ACAAATCATAATTGTTTCAGTAAAATGGTTACTTGGTTTTTCCAAGAAAAAACTCGTTTT 240
170FI       ACAAATCATAATTGTTTCAGTAAAATGGTTACTTGGTTTTTCCAAGAAAAAACTCGTTTT 240
170FII      ACAAATCATAATTGTTTCAGTAAAATGGTTACTTGGTTTTTCCAAGAAAAAACTCGTTTT 240
324RI       ------------------------------------------------------------ 240
324RII      ------------------------------------------------------------ 240
324FI       ------------------------------------------------------------ 240
324FII      ------------------------------------------------------------ 240
```

```
170RI       ------------------------------------------------------------ 240

170RII      ------------------------------------------------------------ 240

expected    TACTCATTTTTGGCCTGTTTGTTTATTTAGAAACTAATCTGGATTCACTCCCTCTGGTTG 300
genomic     TACTCATTTTTGGCCTGTTTGTTTATTTAGAAACTAATCTGGATTCACTCCCTCTGGTTG 300
170FI       TACTCATTTTTGGCCTGTTTGTTTATTTAGAAACTAATCTGGATTCACTCCCTCTGGTTG 300
170FII      TACTCATTTTTGGCCTGTTTGTTTATTTAGAAACTAATCTGGATTCACTCCCTCTGGTTG 300
324RI       ------------------------------------------------------------ 300
324RII      ------------------------------------------------------------ 300
324FI       ------------------------------------------------------------ 300
324FII      ------------------------------------------------------------ 300
170RI       ------------------------------------------------------------ 300
170RII      ------------------------------------------------------------ 300

expected    ATACCCACTCAAAAAGGACACTTCTGATTAAGACGGTTGAAACTAGAGATGGACAGGTTG 360
genomic     ATACCCACTCAAAAAGGACACTTCTGATTAAGACGGTTGAAACTAGAGATGGACAGGTTG 360
170FI       ATACCCACTCAAAAAGGACACTTCTGATTAAGACGGTTGAAACTAGAGATGGACAGGTTG 360
170FII      ATACCCACTCAAAAAGGACACTTCTGATTAAGACGGTTGAAACTAGAGATGGACAGGTTG 360
324RI       ------------------------------------------------------------ 360
324RII      ------------------------------------------------------------ 360
324FI       ------------------------------------------------------------ 360
324FII      ------------------------------------------------------------ 360
170RI       ------------------------------------------------------------ 360
170RII      ------------------------------------------------------------ 360

expected    GTATCTTTTAAGGAAAAAAATAGGGTAATCTCAGACAGGAGTTGATATATTTTAAAATCAG 420
genomic     GTATCTTTTAAGGAAAAAAATAGGGTAATCTCAGACAGGAGTTGATATATTTTAAAATCAG 420
170FI       GTATCTTTTAAGGAAAAAAATAGGGTAATCTCAGACAGGAGTTGATATATTTTAAAATCAG 420
170FII      GTATCTTTTAAGGAAAAAAATAGGGTAATCTCAGACAGGAGTTGATATATTTTAAAATCAG 420
324RI       ----------------------------------------------------AAAATCAG 420
324RII      ----------------------------------------------------AAAATCAG 420
324FI       ------------------------------------------------------------ 420
324FII      ------------------------------------------------------------ 420
170RI       ------------------------------------------------------------ 420
170RII      ------------------------------------------------------------ 420

expected    TGAATCTGAATCTCAGATACAGCTGGCTAATTTGAGAGGTTCAGGTTTCATTCATGCCTA 480
genomic     TGAATCTGAATCTCAGATACAGCTGGCTAATTTGAGAGGTTCAGGTTTCATTCATGCCTA 480
170FI       TGAATCTGAATCTCAGATACAGCTGGCTAATTTGAGAGGTTCAGGTTTCATTCATGCCTA 480
170FII      TGAATCTGAATCTCAGATACAGCTGGCTAATTTGAGAGGTTCAGGTTTCATTCATGCCTA 480
324RI       TGAATCTGAATCTCAGATACAGCTGGCTAATTTGAGAGGTTCAGGTTTCATTCATGCCTA 480
324RII      TGAATCTGAATTTCAGATACAGCTGGCTAATTTGAGAGGTTCAGGTTTCATTCATGCCTA 480
324FI       ------------------------------------------------------------ 480
324FII      ------------------------------------------------------------ 480
170RI       ------------------------------------------------------------ 480
```

```
170RII      ------------------------------------------------------------ 480

expected    CTAAAAAAAGAATAGGC-TTCTTCTTCCAGCAGTACACACAGCCAACTAATTATTTGGCT 540
genomic     CTAAAAAAAGAATAGGC-TTCTTCTTCCAGCAGTACACACAGCCAACTAATTATTTGGCT 540
170FI       CTAAAAAAAGAATAGGC-TTCTTCTTCCAGCAGTACACACAGCCAACTAATTATTTGGCT 540
170FII      CTAAAAAAAGAATAGGC-TTCTTCTTCCAGCAGTACACACAGCCAACTAATTATTTGGCT 540
324RI       CTAAAAAAAGAATAGGCTTTTTTTTTCCAGCAGTACACACAGCCAACTAATTATTTGGCT 540
324RII      CTAAAAAAAGAATAGGC-TTCTTTTTCCAGCAGTACACACAGCCAACTAATTATTTGGCT 540
324FI       ------------------------------------------------------------ 540
324FII      ------------------------------------------------------------ 540
170RI       ------------------------------------------------------------ 540
170RII      ------------------------------------------------------------ 540

expected    CCTGGATGTGAAGTTGAGATAGCAGTCTTCCTGTGCTCCAGAATTAGTGATTTGCTTTGG 600
genomic     CCTGGATGTGAAGTTGAGATAGCAGTCTTCCTGTGCTCCAGAATTAGTGATTTGCTTTGG 600
170FI       CCTGGATGTGAAGTTGAGATAGCAGTCTTCCTGTGCTCCAGAATTAGTGATTTGCTTTGG 600
170FII      CCTGGATGTGAAGTTGAGATAGCAGTCTTCCTGTGCTCCAGAATTAGTGATTTGCTTTGG 600
324RI       CCTGGATGTGAAGTTGAGATAGCAGTCTTCCTGTGCTCCAGAATTAGTGATTTGCTTTGG 600
324RII      CCTGGATGTGAAGTTGAGATAGCAGTCTTCCTGTGCTCCAGAATTAGTGATTTGCTTTGG 600
324FI       ------------------------------------------------------------ 600
324FII      ------------------------------------------------------------ 600
170RI       ------------------------------------------------------------ 600
170RII      ------------------------------------------------------------ 600

expected    TGCTTAATTTGAAGTGGGAGTAAGCTTCCTTAAACCACTTCCTAAAGCAGCTACATGAAA 660
genomic     TGCTTAATTTGAAGTGGGAGTAAGCTTCCTTAAACCACTTCCTAAAGCAGCTACATGAAA 660
170FI       TGCTTAATTTGAAGTGGGAGTAAGCTTCCTTAAACCACTTCCTAAAGCAGCTACATGAAA 660
170FII      TGCTTAATTTGAAGTGGGAGTAAGCTTCCTTAAACCACTTCCTAAAGCAGCTACATGAAA 660
324RI       TGCTTAATTTGAAGTGGGAGTAAGCTTCCTTAAACCACTTCCTAAAGCAGCTACATGAAA 660
324RII      TGCTTAATTTGAAGTGGGAGTAAGCTTCCTTAAACCACTTCCTAAAGCAGCTACATGAAA 660
324FI       ------------------------------------------------------------ 660
324FII      ------------------------------------------------------------ 660
170RI       ------------------------------------------------------------ 660
170RII      ------------------------------------------------------------ 660

expected    CAGCTTCACTAGACTACCTCAATATGAGGAATGTTTTGATCCTGGACATATGGTGTCTTC 720
genomic     CAGCTTCACTAGACTACCTCAATATGAGGAATGTTTTGATCCTGGACATATGGTGTCTTC 720
170FI       CAGCTTCACTAGACTACCTCAATATGAGGAATGTTTTGATCCTGGACATATGGTGTCTTC 720
170FII      CAGCTTCACTAGACTACCTCAATATGAGGAATGTTTTGATCCTGGACATATGGTGTCTTC 720
324RI       CAGCTTCACTAGACTACCTCAATATGAGGAATGTTTTGATCCTGGACATATGGTGTCTTC 720
324RII      CAGCTTCACTAGACTACCTCAATATGAGGAATGTTTTGATCCTGGACATATGGTGTCTTC 720
324FI       ------------------------------------------------------------ 720
324FII      ------------------------------------------------------------ 720
170RI       ------------------------------------------------------------ 720
170RII      ------------------------------------------------------------ 720
```

```
expected    CTACCTCCATACTTTATAGATTCCTAAACCCATCTATATAATACAAGCATGTGCCATACG 780
genomic     CTACCTCCATACTTTATAGATTCCTAAACCCATCTATATAATACAAGCATGTGCCATACG 780
170FI       CTACCTCCATACTTTATAGATTCCTAAACCCATCTATATAATACAAGCATGTGCCATACG 780
170FII      CTACCTCCATACTTTATAGATTCCTAAACCCATCTATATAATACAAGCATGTGCCATACG 780
324RI       CTACCTCCATACTTTATAGATTCCTAAACCCATCTATATAATACAAGCATGTGCCATACG 780
324RII      CTACCTCCATACTTTATAGATTCCTAAACCCATCTATATAATACAAGCATGTGCCATACG 780
324FI       ------------------------------------------------------------ 780
324FII      ------------------------------------------------------------ 780
170RI       ------------------------------------------------------------ 780
170RII      ------------------------------------------------------------ 780

expected    ATCATTTAGTTTCTTATTACCTCCCTATGCCAGGAAAGAAATAGTTGCAATTTATTGTAG 840
genomic     ATCATTTAGTTTCTTATTACCTCCCTATGCCAGGAAAGAAATAGTTGCAATTTATTGTAG 840
170FI       ATCATTTAGTTTCTTATTACCTCCCTATGCCAGGAAAGAAATAGTTGCAATTTATTGTAG 840
170FII      ATCATTTAGTTTCTTATTACCTCCCTATGCCAGGAAAGAAATAGTTGCAATTTATTGTAG 840
324RI       ATCATTTAGTTTCTTATTACCTCCCTATGCCAGGAAAGAAATAGTTGCAATTTATTGTAG 840
324RII      ATCATTTAGTTTCTTATTACCTCCCTATGCCAGGAAAGAAATAGTTGCAATTTATTGTAG 840
324FI       ------------------------------------------------------------ 840
324FII      ------------------------------------------------------------ 840
170RI       ------------------------------------------------------------ 840
170RII      ------------------------------------------------------------ 840

expected    TCATCATGAAATCTTCCCTTGCACATAAATTTAAAATGTACCTGCTGCACATTTTAATAT 900
genomic     TCATCATGAAATCTTCCCTTGCACATAAATTTAAAATGTACCTGCTGCACATTTTAATAT 900
170FI       TCATCATGAAATCTTCCCTTGCACATAAATTTAAAATGTACCTGCTGCACATTTTAATAT 900
170FII      TCATCATGAAATCTTCCCTTGCACATAAATTTAAAATGTACCTGCTGCACATTTTAATAT 900
324RI       TCATCATGAAATCTTCCCTTGCACATAAATTTAAAATGTACCTGCTGCACATTTTAATAT 900
324RII      TCATCATGAAATCTTCCCTTGCACATAAATTTAAAATGTACCTGCTGCACATTTTAATAT 900
324FI       ------------------------------------------------------------ 900
324FII      ------------------------------------------------------------ 900
170RI       ------------------------------------------------------------ 900
170RII      ------------------------------------------------------------ 900

expected    GTCTTAATTGCTTTTAAACTTGGCTGTATTGTGTACAACTATTATACCATCTTTTATAAA 960
genomic     GTCTTAATTGCTTTTAAACTTGGCTGTATTGTGTACAACTATTATACCATCTTTTATAAA 960
170FI       GTCTTAATTGCTTTTAAACTTGGCTGTATTGTGTACAACTATTATACCATCTTTTATAAA 960
170FII      GTCTTAATTGCTTTTAAACTTGGCTGTATTGTGTACAACTATTATACCATCTTTTATAAA 960
324RI       GTTTTAATTGCTTTTAAACTTGGCTGTATTGTGTACAACTATTATACCATCTTTTATAAA 960
324RII      GTTTTAATTGCTTTTAAACTTGGCTGTATTGTGTACAACTATTATACCATCTTTTATAAA 960
324FI       ------------------------------------------------------------ 960
324FII      ------------------------------------------------------------ 960
170RI       ------------------------------------------------------------ 960
170RII      ------------------------------------------------------------ 960
```

```
expected    CACAGTTTTTTAAGAAATTTCTTTTTGTAAGTTACAACATTCCACTGGAtccttatAGGA 1020
genomic     CACAGTTTTTTAAGAAATTTCTTTTTGTAAGTTACAACATTCCACTGGATCCTTATATTG 1020
170FI       CACAGTTTTTTAAGAAATTTCTTTTTGTAAGTTACAACATTCCACTGGATCCTTATAGGA 1020
170FII      CACAGTTTTTTAAGAAATTTCTTTTTGTAAGTTACAACATTCCACTGGATCCTTATAGGA 1020
324RI       CMCAGTTTTTTWAGAAATTTCTTTTTGTAAGTTACAACMTTYCACTGGATCCTTATWGGA 1020
324RII      CACAGTTTTTTWAGAAATTTCTTTTTGTAAGTTACAACMTTYCACYGGATCCTTATWGGA 1020
324FI       ------------------------------------------------------------ 1020
324FII      ------------------------------------------------------------ 1020
170RI       ------------------------------------------------------------ 1020
170RII      ------------------------------------------------------------ 1020

expected    CATATAATACAAGAGGGTCTTGTGTGTCTGCCCCTTCTAGTTTTCACTCATGCAGAAGCA 1080
genomic     CCT--------------------------------------------------------- 1080
170FI       CATATAATACAAGAGGGTCTTGTGTGTCTGCCCCTTCTAGTTTTCACTCATGCAGAAGCA 1080
170FII      CATATAATACAAGAGGGTCTTGTGTGTCTGCCCCTTCTAGTTTTCACTCATGCAGAAGCA 1080
324RI       CATATAATACMAGAGGGTCTTGKGKGTCTGCCCCTTCTAGTTTTCAMTCATGCAGAAGCA 1080
324RII      CATATAATACMAGAGGGTCTTGKGKGTCTGCCCCTTCTAGTTTTCAMTCATGCAGAAGCA 1080
324FI       ------------------------------------------------------------ 1080
324FII      ------------------------------------------------------------ 1080
170RI       ------------------------------------------------------------ 1080
170RII      ------------------------------------------------------------ 1080

expected    ACATAACCTTCTGATTTGCACAATAAATTACATATATTTAGCAGGATTTTTATTTGCCGT 1140
genomic     ------------------------------------------------------------ 1140
170FI       ACATAACCTTCTGATTTGC----------------------------------------- 1140
170FII      ACATAACCTTCTGATTT------------------------------------------- 1140
324RI       ACATAACCTTYTGATTTGCACAATAAATTACATATATTTAGCMGGATTTTTWTTTGCCST 1140
324RII      ACATAACCTTYTGATTTGCACAATAAATTACAWATATTTAGCMGGATTTTTATTTGCCST 1140
324FI       ------------------------------------------------------------ 1140
324FII      ------------------------------------------------------------ 1140
170RI       ------------------------------------------------------------ 1140
170RII      ------------------------------------------------------------ 1140

expected    GATATATAGGATAATTTAGTCTTTGGCATGTGGCATTATATTTATTTTGGTTTTTTTTTT 1200
genomic     ------------------------------------------------------------ 1200
170FI       ------------------------------------------------------------ 1200
170FII      ------------------------------------------------------------ 1200
324RI       GATATATAGGATAAWTTAGTYTTTGGCMTGKGGCMTTATATTTATTTTGGTTTTTTTTTT 1200
324RII      GATATATAGGATAAWTTAGTYTTTGGCMTGTGGCMTTATATTTATTTTGGTTTTTTTTTT 1200
324FI       ------------------------------------------------------------ 1200
324FII      ------------------------------------------------------------ 1200
170RI       ------------------------------------------------------------ 1200
170RII      ------------------------------------------------------------ 1200
```

```
expected   TAAACAGGTTATCAACGAAACTTCTCAGCATCACGATGACCTTGAAggaagcggagaggg 1260
genomic    ------------------------------------------------------------ 1260
170FI      ------------------------------------------------------------ 1260
170FII     ------------------------------------------------------------ 1260
324RI      TAAACAGGTTATCAACGAAACTTCTYAGCATCACGATGACCTTGAAGGAAGCGGAGAGGG 1260
324RII     TAAACAGGTTATCAACGAAACTTCTYAGCATCACGATGACCYTGAAGGAAGCGGAGAGGG 1260
324FI      ------------------------------------------------------------ 1260
324FII     ------------------------------------------------------------ 1260
170RI      ------------------------------------------------------------ 1260
170RII     ------------------------------------------------------------ 1260

expected   cagaggaagtctgctaacatgcggtgacgtcgaggagaatcctggacctATGGTGAGCAA 1320
genomic    ------------------------------------------------------------ 1320
170FI      ------------------------------------------------------------ 1320
170FII     ------------------------------------------------------------ 1320
324RI      CAGRGGAAGTCTGSTAACMTGCGGTGACGTCGAGGAGAATCCTGGACCTATGGTGAGCAA 1320
324RII     CAGRGGAAGTCTGSTAACMTGCGGTGACGTCGAGGAGAATCCTGGACCTATGGTGAGCAA 1320
324FI      ------------------------------------------------------------ 1320
324FII     ------------------------------------------------------------ 1320
170RI      ------------------------------------------------------------ 1320
170RII     ------------------------------------------------------------ 1320

expected   GGGCGAGGAGCTGATCAAGGAGAACATGCACATGAAGCTGTACATGGAAGGCACCGTGAA 1380
genomic    ------------------------------------------------------------ 1380
170FI      ------------------------------------------------------------ 1380
170FII     ------------------------------------------------------------ 1380
324RI      GGGCGAGGAGCTGATCAAGGAGAACATGCACATGAAGCTGTACATGGAAGGC-------- 1380
324RII     GGGCGAGGAGCTGATCAAGGAGAACATGCACATGAAGCTGTACATGGAAGGCACC----- 1380
324FI      ------------------------------------------------------------ 1380
324FII     ------------------------------------------------------------ 1380
170RI      ------------------------------------------------------------ 1380
170RII     ------------------------------------------------------------ 1380

expected   CAACCACCACTTCAAGTGCACCACCGAAGGGGAGGGCAAGCCCTACGAGGGCACCCAGAC 1440
genomic    ------------------------------------------------------------ 1440
170FI      ------------------------------------------------------------ 1440
170FII     ------------------------------------------------------------ 1440
324RI      ------------------------------------------------------------ 1440
324RII     ------------------------------------------------------------ 1440
324FI      -------------------------------GAGGGCAAGCCCTACGAGGGCACCCAGAC 1440
324FII     ---------------------------------GGGCAAGCCCTACGAGGGCACCCAGAC 1440
170RI      ------------------------------------------------------------ 1440
170RII     ------------------------------------------------------------ 1440

expected   CCAGAGGATTAAGGTGGTGGAGGGAGGCCCCCTGCCGTTCGCATTCGACATCCTGGCCAC 1500
```

```
genomic     ------------------------------------------------------------ 1500
170FI       ------------------------------------------------------------ 1500
170FII      ------------------------------------------------------------ 1500
324RI       ------------------------------------------------------------ 1500
324RII      ------------------------------------------------------------ 1500
324FI       CCAGAGGATTAAGGTGGTGGAGGGAGGCCCCCTGCCGTTCGCATTCGACATCCTGGCCAC 1500
324FII      CCAGAGGATTAAGGTGGTGGAGGGAGGCCCCCTGCCGTTCGCATTCGACATCCTGGCCAC 1500
170RI       ------------------------------------------------------------ 1500
170RII      ------------------------------------------------------------ 1500

expected    CTGCTTTATGTACGGGAGCAAGACCTTCATCAACCACACCCAGGGCATCCCCGATTTCTT 1560
genomic     ------------------------------------------------------------ 1560
170FI       ------------------------------------------------------------ 1560
170FII      ------------------------------------------------------------ 1560
324RI       ------------------------------------------------------------ 1560
324RII      ------------------------------------------------------------ 1560
324FI       CTGCTTTATGTACGGGAGCAAGACCTTCATCAACCACACCCAGGGCATCCCCGATTTCTT 1560
324FII      CTGCTTTATGTACGGGAGCAAGACCTTCATCAACCACACCCAGGGCATCCCCGATTTCTT 1560
170RI       ------------------------------------------------------------ 1560
170RII      ------------------------------------------------------------ 1560

expected    TAAGCAGTCCTTCCCTGAGGGCTTCACATGGGAGAGAGTCACCACATACGAAGACGGGGG 1620
genomic     ------------------------------------------------------------ 1620
170FI       ------------------------------------------------------------ 1620
170FII      ------------------------------------------------------------ 1620
324RI       ------------------------------------------------------------ 1620
324RII      ------------------------------------------------------------ 1620
324FI       TAAGCAGTCCTTCCCTGAGGGCTTCACATGGGAGAGAGTCACCACATACGAAGACGGGGG 1620
324FII      TAAGCAGTCCTTCCCTGAGGGCTTCACATGGGAGAGAGTCACCACATACGAAGACGGGGG 1620
170RI       ------------------------------------------------------------ 1620
170RII      ------------------------------------------------------------ 1620

expected    CGTGCTTACCGTTACCCAGGACACCAGCCTCCAGGACGGCTGCTTGATCTACAACGTCAA 1680
genomic     ------------------------------------------------------------ 1680
170FI       ------------------------------------------------------------ 1680
170FII      ------------------------------------------------------------ 1680
324RI       ------------------------------------------------------------ 1680
324RII      ------------------------------------------------------------ 1680
324FI       CGTGCTTACCGTTACCCAGGACACCAGCCTCCAGGACGGCTGCTTGATCTACAACGTCAA 1680
324FII      CGTGCTTACCGTTACCCAGGACACCAGCCTCCAGGACGGCTGCTTGATCTACAACGTCAA 1680
170RI       ------------------------------------------------------------ 1680
170RII      ------------------------------------------------------------ 1680

expected    GCTCAGAGGGGTGAACTTCCCATCCAACGGCCCTGTGATGCAGAAGAAAACACTCGGCTG 1740
genomic     ------------------------------------------------------------ 1740
```

```
170FI       ------------------------------------------------------------ 1740
170FII      ------------------------------------------------------------ 1740
324RI       ------------------------------------------------------------ 1740
324RII      ------------------------------------------------------------ 1740
324FI       GCTCAGAGGGGTGAACTTCCCATCCAACGGCCCTGTGATGCAGAAGAAAACACTCGGCTG 1740
324FII      GCTCAGAGGGGTGAACTTCCCATCCAACGGCCCTGTGATGCAGAAGAAAACACTCGGCTG 1740
170RI       ------------------------------------------------------------ 1740
170RII      ------------------------------------------------------------ 1740

expected    GGAGGCCACCACCGAGACCCTGTACCCCGCTGACGGCGGCCTGGAAGGCAGATGCGACAT 1800
genomic     ------------------------------------------------------------ 1800
170FI       ------------------------------------------------------------ 1800
170FII      ------------------------------------------------------------ 1800
324RI       ------------------------------------------------------------ 1800
324RII      ------------------------------------------------------------ 1800
324FI       GGAGGCCACCACCGAGACCCTGTACCCCGCTGACGGCGGCCTGGAAGGCAGATGCGACAT 1800
324FII      GGAGGCCACCACCGAGACCCTGTACCCCGCTGACGGCGGCCTGGAAGGCAGATGCGACAT 1800
170RI       ------------------------------------------------------------ 1800
170RII      ------------------------------------------------------------ 1800

expected    GGCCCTGAAGCTCGTGGGCGGGGGCCACCTGCACTGCAACCTGAAGACCACATACAGATC 1860
genomic     ------------------------------------------------------------ 1860
170FI       ------------------------------------------------------------ 1860
170FII      ------------------------------------------------------------ 1860
324RI       ------------------------------------------------------------ 1860
324RII      ------------------------------------------------------------ 1860
324FI       GGCCCTGAAGCTCGTGGGCGGGGGCCACCTGCACTGCAACCTGAAGACCACATACAGATC 1860
324FII      GGCCCTGAAGCTCGTGGGCGGGGGCCACCTGCACTGCAACCTGAAGACCACATACAGATC 1860
170RI       ------------------------------------------------------------ 1860
170RII      ------------------------------------------------------------ 1860

expected    CAAGAAACCCGCTAAGAACCTCAAGATGCCCGGCGTCTACTTTGTGGACCGCAGACTGGA 1920
genomic     ------------------------------------------------------------ 1920
170FI       ------------------------------------------------------------ 1920
170FII      ------------------------------------------------------------ 1920
324RI       ------------------------------------------------------------ 1920
324RII      ------------------------------------------------------------ 1920
324FI       CAAGAAACCCGCTAAGAACCTCAAGATGCCCGGCGTCTACTTTGTGGACCGCAGACTGGA 1920
324FII      CAAGAAACCCGCTAAGAACCTCAAGATGCCCGGCGTCTACTTTGTGGACCGCAGACTGGA 1920
170RI       ------------------------------------------------------------ 1920
170RII      ------------------------------------------------------------ 1920

expected    AAGAATCAAGGAGGCCGACAATGAGACCTACGTCGAGCAGCACGAGGTGGCTGTGGCCAG 1980
genomic     ------------------------------------------------------------ 1980
170FI       ------------------------------------------------------------ 1980
```

```
170FII      ------------------------------------------------------------ 1980
324RI       ------------------------------------------------------------ 1980
324RII      ------------------------------------------------------------ 1980
324FI       AAGAATCAAGGAGGCCGACAATGAGACCTACGTCGAGCAGCACGAGGTGGCTGTGGCCAG 1980
324FII      AAGAATCAAGGAGGCCGACAATGAGACCTACGTCGAGCAGCACGAGGTGGCTGTGGCCAG 1980
170RI       ------------------------------------------------------------ 1980
170RII      ------------------------------------------------------------ 1980

expected    ATACTGCGACCTCCCTAGCAAACTGGGGCACAAACTTAATGGCATGGACGAGCTGTACAA 2040
genomic     ------------------------------------------------------------ 2040
170FI       ------------------------------------------------------------ 2040
170FII      ------------------------------------------------------------ 2040
324RI       ------------------------------------------------------------ 2040
324RII      ------------------------------------------------------------ 2040
324FI       ATACTGCGACCTCCCTAGCAAACTGGGGCACAAACTTAATGGCATGGACGAGCTGTACAA 2040
324FII      ATACTGCGACCTCCCTAGCAAACTGGGGCACAAACTTAATGGCATGGACGAGCTGTACAA 2040
170RI       ------------------------------------------------------------ 2040
170RII      ------------------------------------------------------------ 2040

expected    GGACTATAAAGACCACGACGGAGACTACAAGGATCATGATATTGATTACAAAGACGATGA 2100
genomic     ------------------------------------------------------------ 2100
170FI       ------------------------------------------------------------ 2100
170FII      ------------------------------------------------------------ 2100
324RI       ------------------------------------------------------------ 2100
324RII      ------------------------------------------------------------ 2100
324FI       GGACTATAAAGACCACGACGGAGACTACAAGGATCATGATATTGATTACAAAGACGATGA 2100
324FII      GGACTATAAAGACCACGACGGAGACTACAAGGATCATGATATTGATTACAAAGACGATGA 2100
170RI       ----------GACCACGACGGAGACTACAAGGATCATGATATTGATTACAAAGACGATGA 2100
170RII      -------------------------------------------------AAAGACGATGA 2100

expected    CGATAAGTAATAGTGGAAGAGGGTCTTGTGTGTCTGCCCCTTCTAGTTTTCACTCATGCA 2160
genomic     ---------GTAGTGGAAGAGGGTCTTGTGTGTCTGCCCCTTCTAGTTTTCACTCATGCA 2160
170FI       ------------------------------------------------------------ 2160
170FII      ------------------------------------------------------------ 2160
324RI       ------------------------------------------------------------ 2160
324RII      ------------------------------------------------------------ 2160
324FI       CGATAAGTAATAGTGGAAGAGGGTCTTGTGTGTCTGCCCCTTCTAGTTTTCACTCATGCA 2160
324FII      CGATAAGTAATAGTGGAAGAGGGTCTTGTGTGTCTGCCCCTTCTAGTTTTCACTCATGCA 2160
170RI       CGATAAGTAATAGTGGAAGAGGGTCTTGTGTGTCTGCCCCTTCTAGTTTTCACTCATGCA 2160
170RII      CGATAAGTTATAGTGGAAGAGGGTCTTGTGTGTCTGCCCCTTCTAGTTTTCACTCATGCA 2160

expected    GAAGCAACATAACCTTCTGATTTGCACAATAAATTACATATATTTAGCAGGATTTTTATT 2220
genomic     GAAGCAACATAACCTTCTGATTTGCACAATAAATTACATATATTTAGCAGGATTTTTATT 2220
170FI       ------------------------------------------------------------ 2220
170FII      ------------------------------------------------------------ 2220
```

```
324RI       ------------------------------------------------------------ 2220
324RII      ------------------------------------------------------------ 2220
324FI       GAAGCAACATAACCTTCTGATTTGCACAATAAATTACATATATTTAGCAGGATTTTTATT 2220
324FII      GAAGCAACATAACCTTCTGATTTGCACAATAAATTACATATATTTAGCAGGATTTTTATT 2220
170RI       GAAGCAACATAACCTTCTGATTTGCACAATAAATTACATATATTTAGCAGGATTTTTATT 2220
170RII      GAAGCAACATAACCTTCTGATTTGCACAATAAATTACATATATTTAGCAGGATTTTTATT 2220

expected    TGCCGTGATATATAGGATAATTTAGTCTTTGGCATGTGGCATTATATTTATTTTGGTTTT 2280
genomic     TGCCGTGATATATAGGATAATTTAGTCTTTGGCATGTGGCATTATATTTATTTTGGTTTT 2280
170FI       ------------------------------------------------------------ 2280
170FII      ------------------------------------------------------------ 2280
324RI       ------------------------------------------------------------ 2280
324RII      ------------------------------------------------------------ 2280
324FI       TGCCGTGATATATAGGATAATTTAGTCTTTGGCATGTGGCATTATATTTATTTTGG-TTT 2280
324FII      TGCCGTGAtATATAGGATAATTTAGTCTTTGGCATGTGGCATTATATTTATTTTGG-TTT 2280
170RI       TGCCGTGATATATAGGATAATTTAGTCTTTGGCATGTGGCATTATATTTATTTTGGTTTT 2280
170RII      TGCCGTGATATATAGGATAATTTAGTCTTTGGCATGTGGCATTATATTTATTTTGGTTTT 2280

expected    TTTTTTTAAACAGGTTATCAACGAAACTTCTCAGCATCACGATGACCTTGAATAAAAATT 2340
genomic     TTTTTTTAAACAGGTTATCAACGAAACTTCTCAGCATCACGATGACCTTGAATAAAAATT 2340
170FI       ------------------------------------------------------------ 2340
170FII      ------------------------------------------------------------ 2340
324RI       ------------------------------------------------------------ 2340
324RII      ------------------------------------------------------------ 2340
324FI       TTTTTTTAAACAGGTTATCAACGAAACTTCTCAGCATCACGATGACCTTGAATAAAAATT 2340
324FII      TTTTTTTAAACAGGTTATCAACGAAACTTCTCAGCATCACGATGACCTTGAATAAAAATT 2340
170RI       TTTTTTTAAACAGGTTATCAACGAAACTTCTCAGCATCACGATGACCTTGAATAAAAATT 2340
170RII      TTTTTTTAAACAGGTTATCAACGAAACTTCTCAGCATCACGATGACCTTGAATAAAAATT 2340

expected    GCACACACTCAGTGCAGCAATATATTACCAGCAAGAATAAAAAAGAAATCCATATCTTAA 2400
genomic     GCACACACTCAGTGCAGCAATATATTACCAGCAAGAATAAAAAAGAAATCCATATCTTAA 2400
170FI       ------------------------------------------------------------ 2400
170FII      ------------------------------------------------------------ 2400
324RI       ------------------------------------------------------------ 2400
324RII      ------------------------------------------------------------ 2400
324FI       GCAC-------------------------------------------------------- 2400
324FII      GCACACACTCAGTGCAGCAATATATTACCAGCAAGA--TAAAAAGAAATCCATATC-TAA 2400
170RI       GCACACACTCAGTGCAGCAATATATTACCAGCAAGAATAAAAAAGAAATCCATATCTTAA 2400
170RII      GCACACACTCAGTGCAGCAATATATTACCAGCAAGAATAAAAAAGAAATCCATATCTTAA 2400

expected    AGAAACAGCTTTCAAGTGCCTTTCTGCAGTTTTTCAGGAGCGCAAGATAGATTTGGAATA 2460
genomic     AGAAACAGCTTTCAAGTGCCTTTCTGCAGTTTTTCAGGAGCGCAAGATAGATTTGGAATA 2460
170FI       ------------------------------------------------------------ 2460
170FII      ------------------------------------------------------------ 2460
324RI       ------------------------------------------------------------ 2460
```

```
324RII      ------------------------------------------------------------ 2460
324FI       ------------------------------------------------------------ 2460
324FII      AGAAACAGCTTTC-AGTGCCTTTCTGCAGTTTT--------------------------- 2460
170RI       AGAAACAGCTTTCAAGTGCCTTTCTGCAGTTTTTCAGGAGCGCAAGATAGATTTGGAATA 2460
170RII      AGAAACAGCTTTCAAGTGCCTTTCTGCAGTTTTTCAGGAGCGCAAGATAGATTTGGAATA 2460

expected    GGAATAAGCTCTAGTTCTTAACAACCGACACTCCTACAAGATTTAGAAAAAAGTTTACAA 2520
genomic     GGAATAAGCTCTAGTTCTTAACAACCGACACTCCTACAAGATTTAGAAAAAAGTTTACAA 2520
170FI       ------------------------------------------------------------ 2520
170FII      ------------------------------------------------------------ 2520
324RI       ------------------------------------------------------------ 2520
324RII      ------------------------------------------------------------ 2520
324FI       ------------------------------------------------------------ 2520
324FII      ------------------------------------------------------------ 2520
170RI       GGAATAAGCTCTAGTTCTTAACAACCGACACTCCTACAAGATTTAGAAAAAAGTTTACAA 2520
170RII      GGAATAAGCTCTAGTTCTTAACAACCGACACTCCTACAAGATTTAGAAAAAAGTTTACAA 2520

expected    CATAATCTAGTTTACAGAAAAATCTTGTGCTAGAATACTTTTTAAAAGGTATTTTGAATA 2580
genomic     CATAATCTAGTTTACAGAAAAATCTTGTGCTAGAATACTTTTTAAAAGGTATTTTGAATA 2580
170FI       ------------------------------------------------------------ 2580
170FII      ------------------------------------------------------------ 2580
324RI       ------------------------------------------------------------ 2580
324RII      ------------------------------------------------------------ 2580
324FI       ------------------------------------------------------------ 2580
324FII      ------------------------------------------------------------ 2580
170RI       CATAATCTAGTTTACAGAAAAATCTTGTGCTAGAATACTTTTTAAAAGGTATTTTGAATA 2580
170RII      CATAATCTAGTTTACAGAAAAATCTTGTGCTAGAATACTTTTTAAAAGGTATTTTGAATA 2580

expected    CCATTAAAACTGCTTTTTTTTTTCCAGCAAGTATCCAACCAACTTGGTTCTGCTTCAATA 2640
genomic     CCATTAAAACTGCTTTTTTTTTTCCAGCAAGTATCCAACCAACTTGGTTCTGCTTCAATA 2640
170FI       ------------------------------------------------------------ 2640
170FII      ------------------------------------------------------------ 2640
324RI       ------------------------------------------------------------ 2640
324RII      ------------------------------------------------------------ 2640
324FI       ------------------------------------------------------------ 2640
324FII      ------------------------------------------------------------ 2640
170RI       CCATTAAAACTGCTTTTTTTTTTCCAGCAAGTATCCAACCAACTTGGTTCTGCTTCAATA 2640
170RII      CCATTAAAACTGCTTTTTTTTTTCCAGCAAGTATCCAACCAACTTGGTTCTGCTTCAATA 2640

expected    AATCTTTGGAAAAACTCTTTTGTTGTGTTATTTATTGGATAATATCTAAACAATTCTCTA 2700
genomic     AATCTTTGGAAAAACTCTTTTGTTGTGTTATTTATTGGATAATATCTAAACAATTCTCTA 2700
170FI       ------------------------------------------------------------ 2700
170FII      ------------------------------------------------------------ 2700
324RI       ------------------------------------------------------------ 2700
324RII      ------------------------------------------------------------ 2700
```

```
324FI       ------------------------------------------------------------ 2700
324FII      ------------------------------------------------------------ 2700
170RI       AATCTTTGGAAAAACTCTTTTGTTGTGTTATTTATTGGATAATATCTAAACAATTCTCTA 2700
170RII      AATCTTTGGAAAAACTCTTTTGTTGTGTTATTTATTGGATAATATCTAAACAATTCTCTA 2700

expected    CTTGGTCCTATTAGTTAATTTGTCATTACAATCATGTAAGTTGATAAATTCAGGTTATTT 2760
genomic     CTTGGTCCTATTAGTTAATTTGTCATTACAATCATGTAAGTTGATAAATTCAGGTTATTT 2760
170FI       ------------------------------------------------------------ 2760
170FII      ------------------------------------------------------------ 2760
324RI       ------------------------------------------------------------ 2760
324RII      ------------------------------------------------------------ 2760
324FI       ------------------------------------------------------------ 2760
324FII      ------------------------------------------------------------ 2760
170RI       CTTGGTCCTATTAGTTAATTTGTCATTACAATCATGTAAGTTGATAAATTCAGGTTATTT 2760
170RII      CTTGGTCCTATTAGTTAATTTGTCATTACAATCATGTAAGTTGATAAATTCAGGTTATTT 2760

expected    ATGCTTGAGATGTAGTTCTTAATTTTGTCATTTTTGATAGACCTCACTTCTTTTTATTAT 2820
genomic     ATGCTTGAGATGTAGTTCTTAATTTTGTCATTTTTGATAGACCTCACTTCTTTTTATTAT 2820
170FI       ------------------------------------------------------------ 2820
170FII      ------------------------------------------------------------ 2820
324RI       ------------------------------------------------------------ 2820
324RII      ------------------------------------------------------------ 2820
324FI       ------------------------------------------------------------ 2820
324FII      ------------------------------------------------------------ 2820
170RI       ATGCTTGAGATGTAGTTCTTAATTTTGTCATTTTTGATAGACCTCACTTCTTTTTATTAT 2820
170RII      ATGCTTGAGATGTAGTTCTTAATTTTGTCATTTTTGATAGACCTCACTTCTTTTTATTAT 2820

expected    TACTTAAAAACATTTACAAATAGGTGGTGTCGAATAAAATAACTTGTACCAAAATGAAGA 2880
genomic     TACTTAAAAACATTTACAAATAGGTGGTGTCGAATAAAATAACTTGTACCAAAATGAAGA 2880
170FI       ------------------------------------------------------------ 2880
170FII      ------------------------------------------------------------ 2880
324RI       ------------------------------------------------------------ 2880
324RII      ------------------------------------------------------------ 2880
324FI       ------------------------------------------------------------ 2880
324FII      ------------------------------------------------------------ 2880
170RI       TACTTAAAAACATTTACAAATAGGTGGTGTCGAATAAAATAACTTGTACCAAAATGAAGA 2880
170RII      TACTTAAAAACATTTACAAATAGGTGGTGTCGAATAAAATAACTTGTACCAAAATGAAGA 2880

expected    TAGGTCTCTCTAAAATGAGCTCAGGTCTGTGATTTTAATCATAACAAACAGACTTCCTAA 2940
genomic     TAGGTCTCTCTAAAATGAGCTCAGGTCTGTGATTTTAATCATAACAAACAGACTTCCTAA 2940
170FI       ------------------------------------------------------------ 2940
170FII      ------------------------------------------------------------ 2940
324RI       ------------------------------------------------------------ 2940
324RII      ------------------------------------------------------------ 2940
324FI       ------------------------------------------------------------ 2940
```

```
324FII     ------------------------------------------------------------ 2940
170RI      TAGGTCTCTCTAAAATGAGCTCAGGTCTGTGATTTTAATCATAACAAACAGACTTCCTAA 2940
170RII     TAGGTCTCTCTAAAATGAGCTCAGGTCTGTGATTTTAATCATAACAAACAGACTTCCTAA 2940

expected   AATTAAAAAATAAAAACTTTTTTTAGTACATATAGCATATGAGTAACACAAATTCCACTT 3000
genomic    AATTAAAAAATAAAAACTTTTTTTAGTACATATAGCATATGAGTAACACAAATTCCACTT 3000
170FI      ------------------------------------------------------------ 3000
170FII     ------------------------------------------------------------ 3000
324RI      ------------------------------------------------------------ 3000
324RII     ------------------------------------------------------------ 3000
324FI      ------------------------------------------------------------ 3000
324FII     ------------------------------------------------------------ 3000
170RI      AATTAAAAAATAAAAACTTTTTTTAGTACATATAGCATATGAGTAACACA---------- 3000
170RII     AATTAAAAAATAAAAACTTTTTTTAGTACATATAGCATATGAGTAACACAAA-------- 3000

expected   TGGGAGTTAAAGCATAGGAAGTTGCCAAGATATAGGGGCTTATCTTCCGCTAGCAAGATG 3060
genomic    TGGGAGT----------------------------------------------------- 3060
170FI      ------------------------------------------------------------ 3060
170FII     ------------------------------------------------------------ 3060
324RI      ------------------------------------------------------------ 3060
324RII     ------------------------------------------------------------ 3060
324FI      ------------------------------------------------------------ 3060
324FII     ------------------------------------------------------------ 3060
170RI      ------------------------------------------------------------ 3060
170RII     ------------------------------------------------------------ 3060

expected   CAGAGAAATG 3070
genomic    ---------- 3070
170FI      ---------- 3070
170FII     ---------- 3070
324RI      ---------- 3070
324RII     ---------- 3070
324FI      ---------- 3070
324FII     ---------- 3070
170RI      ---------- 3070
170RII     ---------- 3070
```

**Figure S2.** The DNA template for knock-in into *Vimentin* gene. Partial vKiT plasmid sequence which contains the template for the knock-in. The sequence recognized by Cas9/gRNA complex was recoded are shown in red text. Blue text are the PAM sequences.

VIM gRNA target site – 5'HA – VIM missing part - T2A - mCardinal –3xFLAG- 3'HA - VIM gRNA target site

**GATCCTTATATTGCCTGTAGTGGAAGAAAAAACTCGTTTTTACTCATTTTTGGCCTGTTTGTTTATTTAGAAACTAATCT**
**GGATTCACTCCCTCTGGTTGATACCCACTCAAAAAGGACACTTCTGATTAAGACGGTTGAAACTAGAGATGGACAGGTTG**
**GTATCTTTTAAGGAAAAAATAGGGTAATCTCAGACAGGAGTTGATATATTTTAAAATCAGTGAATCTGAATCTCAGATAC**
**AGCTGGCTAATTTGAGAGGTTCAGGTTTCATTCATGCCTACTAAAAAAAGAATAGGCTTCTTCTTCCAGCAGTACACACA**

**GCCAACTAATTATTTGGCTCCTGGATGTGAAGTTGAGATAGCAGTCTTCCTGTGCTCCAGAATTAGTGATTTGCTTTGGT**
**GCTTAATTTGAAGTGGGAGTAAGCTTCCTTAAACCACTTCCTAAAGCAGCTACATGAAACAGCTTCACTAGACTACCTCA**
**ATATGAGGAATGTTTTGATCCTGGACATATGGTGTCTTCCTACCTCCATACTTTATAGATTCCTAAACCCATCTATATAA**
**TACAAGCATGTGCCATACGATCATTTAGTTTCTTATTACCTCCCTATGCCAGGAAAGAAATAGTTGCAATTTATTGTAGT**
**CATCATGAAATCTTCCCTTGCACATAAATTTAAAATGTACCTGCTGCACATTTTAATATGTCTTAATTGCTTTTAAACTT**
**GGCTGTATTGTGTACAACTATTATACCATCTTTTATAAACACAGTTTTTAAGAAATTTCTTTTTGTAAGTTACAACATT**
**CCACTGGA***tccttat***AGGACATATAATAC**AAGAGGGTCTTGTGTGTCTGCCCCTTCTAGTTTTCACTCATGCAGAAGCAA
CATAACCTTCTGATTTGCACAATAAATTACATATATTTAGCAGGATTTTTATTTGCCGTGATATATAGGATAATTTAGTC
TTTGGCATGTGGCATTATATTTATTTTGGTTTTTTTTTTTAAACAGGTTATCAACGAAACTTCTCAGCATCACGATGACC
TTGAAggaagcggagagggcagaggaagtctgctaacatgcggtgacgtcgaggagaatcctggacctATGGTGAGCAAGGGCGAGGAGCTGATCAA
GGAGAACATGCACATGAAGCTGTACATGGAAGGCACCGTGAACAACCACCACTTCAAGTGCACCACCGAAGGGGAGGGCAAGCC
CTACGAGGGCACCCAGACCCAGAGGATTAAGGTGGTGGAGGGAGGCCCCCTGCCGTTCGCATTCGACATCCTGGCCACCTGCTTT
ATGTACGGGAGCAAGACCTTCATCAACCACACCCAGGGCATCCCCGATTTCTTTAAGCAGTCCTTCCCTGAGGGCTTCACATGGGA
GAGAGTCACCACATACGAAGACGGGGGCGTGCTTACCGTTACCCAGGACACCAGCCTCCAGGACGGCTGCTTGATCTACAACGTC
AAGCTCAGAGGGGTGAACTTCCCATCCAACGGCCCTGTGATGCAGAAGAAAACACTCGGCTGGGAGGCCACCACCGAGACCCTGT
ACCCCGCTGACGGCGGCCTGGAAGGCAGATGCGACATGGCCCTGAAGCTCGTGGGCGGGGGCCACCTGCACTGCAACCTGAAGA
CCACATACAGATCCAAGAAACCCGCTAAGAACCTCAAGATGCCCGGCGTCTACTTTGTGGACCGCAGACTGGAAAGAATCAAGGA
GGCCGACAATGAGACCTACGTCGAGCAGCACGAGGTGGCTGTGGCCAGATACTGCGACCTCCCTAGCAAACTGGGGCACAAACTT
AATGGCATGGACGAGCTGTACAAGGACTA**TAAA**GACCACGACGGAGACTACAAGGATCATGATATTGATTACAAAGACG
ATGACGATAAGTAATAGTGGAAGAGGGTCTTGTGTGTCTGCCCCTTCTAGTTTTCACTCATGCAGAAGCAACATAACCTTCTGATT
TGCACAATAAATTACATATATTTAGCAGGATTTTTATTTGCCGTGATATATAGGATAATTTAGTCTTTGGCATGTGGCATTATATTTA
TTTTGGTTTTTTTTTTTAAACAGGTTATCAACGAAACTTCTCAGCATCACGATGACCTTGAATAAAAATTGCACACACTCAGTGCAGC
AATATATTACCAGCAAGAATAAAAAAGAAATCCATATCTTAAAGAAACAGCTTTCAAGTGCCTTTCTGCAGTTTTTCAGGAGCGCA
AGATAGATTTGGAATAGGAATAAGCTCTAGTTCTTAACAACCGACACTCCTACAAGATTTAGAAAAAAGTTTACAACATAATCTAG
TTTACAGAAAAATCTTGTGCTAGAATACTTTTTAAAAGGTATTTTGAATACCATTAAAACTGCTTTTTTTTTTCCAGCAAGTATCCAAC
CAACTTGGTTCTGCTTCAATAAATCTTTGGAAAAACTCTTTTGTTGTGTTATTTATTGGATAATATCTAAACAATTCTCTACTTGGTCC
TATTAGTTAATTTGTCATTACAATCATGTAAGTTGATAAATTCAGGTTATTTATGCTTGAGATGTAGTTCTTAATTTTGTCATTTTTGA
TAGACCTCACTTCTTTTTATTATTACTTAAAAACATTTACAAATAGGTGGTGTCGAATAAAATAACTTGTACCAAAATGAAGATAGG
TCTCTCTAAAATGAGCTCAGGTCTGTCAC**TACAGGCAATATAAGGATC**

**Figure S3.** Confocal imaging of HEK293 cells transfected with re-cloned *VIM-T2A-mCardinal* **(A-C)** and fusion VIM-mCardinal **(D-F).** Cellular distribution of mCardinal in *VIM-T2A-mCardinal* expressing cells shows its strong fluorescence in single dot-like area **(B)**, as well as it is weakly diffused in the cytoplasm. VIM-mCardinal fusion protein aggregated in the cytoplasm **(E)**. **A)** merged, **B)** mCardinal, **C)** transmission light, **D)** merged, **E)** mCardinal, **F)** transmission light. The scale bar is 10 μm.

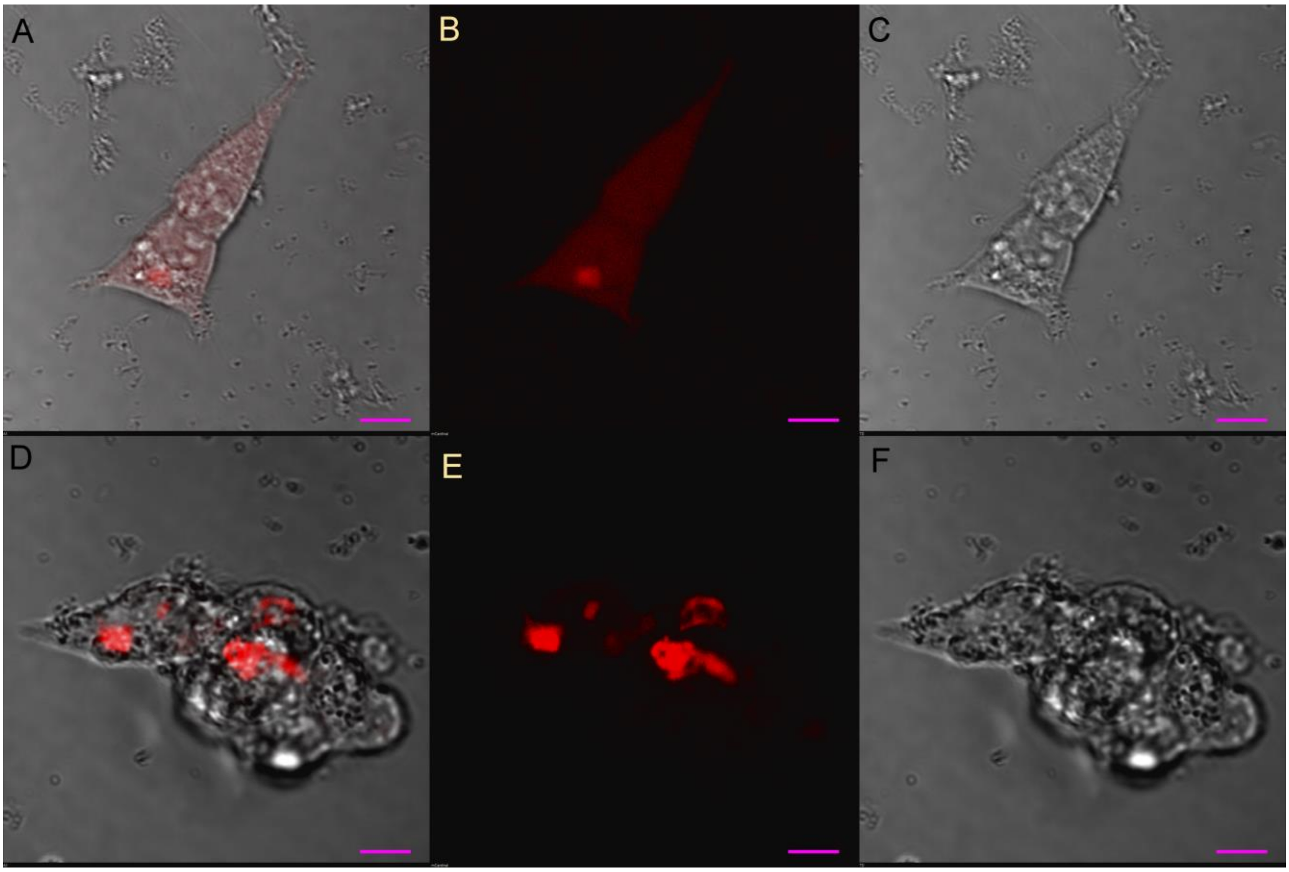

**Figure S4.** Sorting of VRCs by flow cytometry. **A)** P1 acquisition gate was set on FSC and SSC detectors. **B)** the cells showing red fluorescence were selected by P2 or P3 gate, then collected into the tubes and named as dim-VRCs or bright-VRCs respectively. Representative dot-plot has been shown.

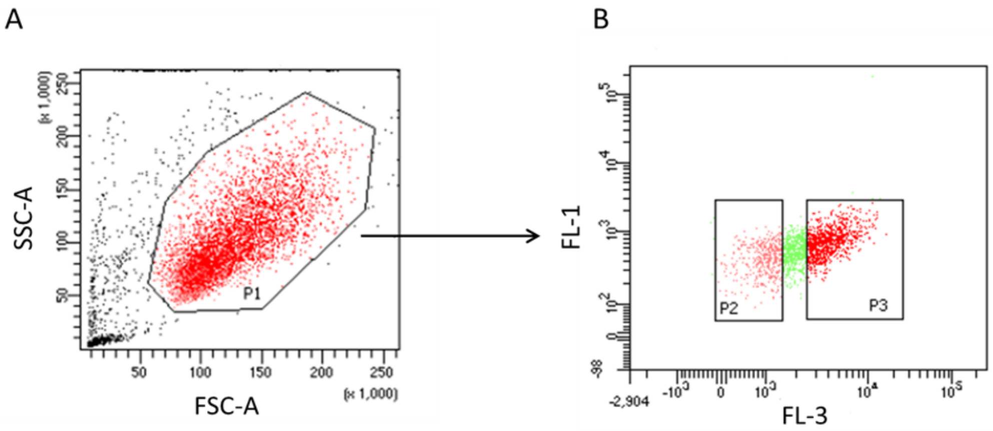


**Figure S5.** Expression of *VIM*, *mCardinal* and *CDH1* in dim-VRCs and bright-VRCs by qPCR. Mean 2^(–ΔCT) ± SEM is shown, the reference gene was *GAPDH*. The graph shows the representative result of the measurement which was done in triplicate.

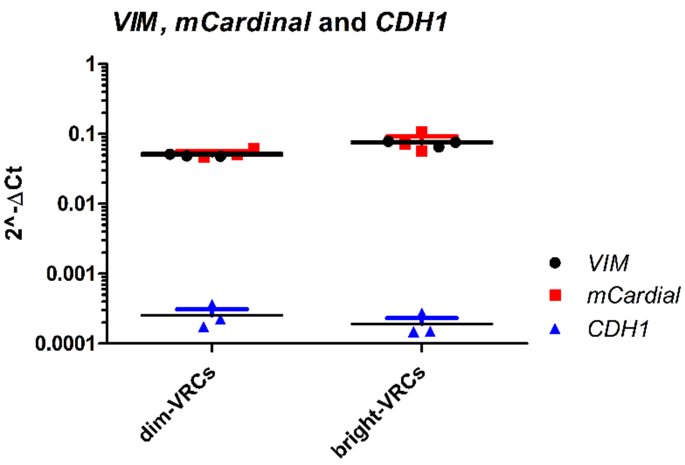


**Figure S6.** EMT markers in VRC and in H2170 parental cells assessed by qPCR. Relative quantification (RQ). Expression of *CDH1*, *VIM*, *SNAI1*, *ZEB1*, *ZEB2*, *TWIST1* and *TWIST2* in VRCs and in parental H2170 cell line has been shown as mean 2^(–ΔΔCT) ± SEM, with the *GAPDH* reference. The graph shows the representative result of the measurement which was done in triplicate.

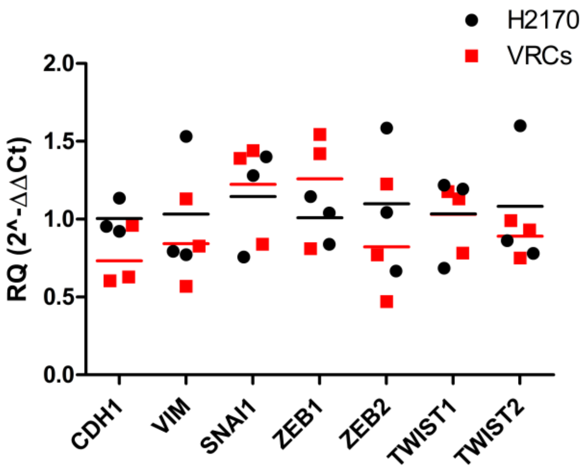

**Table 1.** Oligonucleotides.

| | Name of the oligonucleotide | Sequence |
|---|---|---|
| 1 | gRNA-F | CACCGGATCCTTATATTGCCTGTAG |
| 2 | gRNA-R | AAACCTACAGGCAATATAAGGATCC |
| 3 | KI-F | ATGTCCACCAGGTCCGTGTC |
| 4 | KI-R | CAGACACACAAGACCCTCTTCCAC |
| 5 | BCB-F | AGGGTCTTGTGTGTCTGTCTCAGATCTCGAGCTCAAGCTTC |
| 6 | BCB-R | ACGGACCTGGTGGACATGTACGGAGACGGGATCTGACG |
| 7 | SDM-F | AGGAGAATGGCGGAGGGATGGTGAGCAAGGGCGAGG |
| 8 | SDM-R | CCCTCCGCCATTCTCCTCGACGTCACCGC |
| 9 | 170 | TCGAACCCAAGTACACTCTTGC |
| 10 | 249 | ATCCTTGTAGTCTCCGTCGTGGTC |
| 11 | GAPDH-F | CTCTGCTCCTCCTGTTCGAC |
| 12 | GAPDH-R | GCCCAATACGACCAAATCC |
| 13 | VIM-F | AGTCCACTGAGTACCGGAGAC |
| 14 | VIM-R | CATTTCACGCATCTGGCGTTC |
| 15 | mCard-F | TGATCAAGGAGAACATGCACATGAAGC |
| 16 | mCard-R | CCTTAATCCTCTGGGTCTGGGTG |
| 17 | Cdh1-F | CGAGAGCTACACGTTCACGG |
| 18 | Cdh1-R | GGGTGTCGAGGGAAAAATAGG |
| 19 | Mir145-F | AAGATCTCTACAGATGGGGCTGGATGC |
| 20 | Mir145-R | AAAGCTTCAAGAGTACGGCAGTGCTGA |
| 21 | Mir-200b-F | AAGATCTGGATTAGGACGCTCAGGTGTC |
| 22 | Mir-200b-R | AAAGCTTTGGAGTAGGAGCTCCGGATGTG |
| 23 | Mir-200c-F | AAGATCTAGGGTGGGTAAATCGGTGTG |
| 24 | Mir-200c-R | AAAGCTTACCTGAGGCGATGGATGTTG |
| 25 | Mir-205-F | AAGATCTCCTCCTTGGAGGATGTGA |
| 26 | Mir-205-R | AAAGCTTACGCACACTCCAGATGTCTC |
| 27 | ZEB1F | ACTGTGGTAGAAACAAATTCAGATTCAGATGATG |
| 28 | ZEB1R | GCCCTTCCTTTCCTGTGTCATCC |
| 29 | ZEB2F | CTCTGTAGATGGTCCAGTGAAGAATGC |
| 30 | ZEB2R | GTCACTGCGCTGAAGGTACTCC |
| 31 | SNAI1F | TCGGAAGCCTAACTACAGCGA |
| 32 | SNAI1R | AGATGAGCATTGGCAGCGAG |
| 33 | TWIST1F | GCCGGAGACCTAGATGTCATT |
| 34 | TWIST1R | TTTTAAAAGTGCGCCCCACG |
| 35 | TWIST2F | CGCCAGGGCTGTCCGTC |
| 36 | TWIST2R | GTCACTGCTGTCCCTTCTCTCG |
